# Sex dependent compensatory mechanisms to preserve blood pressure homeostasis in PGI_2_ receptor deficient mice

**DOI:** 10.1101/2020.01.27.921791

**Authors:** Soon Yew Tang, Seán T. Anderson, Hu Meng, Dimitra Sarantopoulou, Emanuela Ricciotti, Elizabeth J. Hennessy, Gregory R. Grant, Garret A. FitzGerald

**Affiliations:** From the Institute for Translational Medicine and Therapeutics, Perelman School of Medicine, Department of Systems Pharmacology and Translational Therapeutics; Department of Genetics, University of Pennsylvania, Philadelphia, Pennsylvania, 19104-5127; National Institute on Aging, National Institutes of Health, 21224.

**Author notes:** Address for correspondence: Garret A. FitzGerald, Institute for Translational Medicine and Therapeutics, Perelman School of Medicine, 10-110 Smilow Center for Translational Research, 3400 Civic Center Blvd, Bldg 421, University of Pennsylvania, Philadelphia, PA 19104-5158.

**Keywords:** Ip, mPges-1, ANP, hyperlipidemic, hypertension

## Abstract

Inhibitors of microsomal prostaglandin E synthase-1 (mPges-1) are in the early phase of clinical development. Deletion of mPges-1 confers analgesia, restrains atherogenesis and fails to accelerate thrombogenesis, while suppressing prostaglandin (PG) E_2_, but increasing biosynthesis of prostacyclin (PGI_2_). In hyperlipidemic mice, this last effect represents the dominant mechanism by which mPges-1 deletion restrains thrombogenesis, while suppression of PGE_2_ accounts for its anti-atherogenic effect. However, the impact of mPges-1 depletion on blood pressure (BP) in this setting remains unknown.

To address how differential effects on PGE_2_ and PGI_2_ might modulate salt-evoked BP responses in the absence of mPges-1, we generated mice lacking the I prostanoid (Ipr) receptor or mPges-1 on a hyperlipidemic background caused by deletion of the low density lipoprotein receptor (Ldlr KOs). Here, mPges-1 depletion significantly increased the BP response to salt loading in male Ldlr KO mice, whereas, despite the direct vasodilator properties of PGI_2_, Ipr deletion suppressed it. Furthermore, combined deletion of the Ipr abrogated the exaggerated BP response in male mPges-1 KO mice. Suppression of PGE_2_ biosynthesis was enough to explain the exaggerated BP response to salt loading by either mPges-1/Ldlr depletion or by an MPGES-1 inhibitor in mice expressing human mPGES-1. However, the lack of a hypertensive response to salt in Ipr-deficient mice was attributable to reactive activation of the atrial natriuretic peptide pathway. Interestingly, these unexpected BP phenotypes were not observed in female mice fed a high salt diet. This is attributable to the protective effect of estrogen in Ldlr KO mice and in Ipr /Ldlr DKOs. Thus, estrogen compensates for a deficiency in PGI_2_ to maintain BP homeostasis in response to high salt in hyperlipidemic female mice. In males, by contrast, augmented formation of ANP plays a similar compensatory role, restraining hypertension and oxidant stress in the setting of Ipr depletion. Hyperlipidemic males on a high salt diet might be at risk of a hypertensive response to mPGES-1 inhibitors.

## Introduction

Both the adverse cardiovascular events associated with non-steroidal anti-inflammatory drugs (NSAIDs) and the opioid crisis have prompted interest in developing new analgesics (1–4). Several clinical trials have shown that the incidence and severity of hypertension from NSAID use is quite variable in humans (5–8). Inhibitors of the microsomal PGE synthase −1 (mPGES-1), an enzyme involved in the biosynthesis of prostaglandin (PG) E_2_, are in early clinical development as potential non-addictive analgesics devoid of the cardiovascular hazards attributable to inhibition of cyclooxygenase-2 (COX-2) by NSAIDs.

Deletion of mPges-1 has a mild adverse cardiovascular profile in normolipidemic mice (3) and we have reported that rediversion of the mPGES-1 substrate prostaglandin (PG)H_2_ to prostacyclin (PGI_2_) synthase, augmenting PGI_2_, attenuates thrombogenesis in hyperlipidemic mice (9). This is a point of distinction from Cox-2 depletion or inhibition that suppresses synthesis of this endogenous platelet inhibitor and predisposes mice to thrombogenic stimuli (3). The sexual dimorphism in blood pressure (BP) homeostasis is at least partly explained by the endocrine system. For example, systolic blood pressure (SBP) is higher in boys from 13 years on compared with girls of the same age (10) and the hypertensive response to salt loading is more pronounced in apparently healthy males compared to premenopausal females at different ages (11). Similarly, in genetically and experimentally predisposed rodent models, hypertension develops more slowly in female than in male mice (12, 13). Deletion of prostaglandin E_2_ receptor, Epr1, reduced BP in male but not female mice (14).

Here, the BP response to a high salt diet (HSD) is augmented in hyperlipidemic mice lacking the low-density lipoprotein receptor (Ldlr). Both PGE_2_ and PGI_2_ may act as direct vasodilators so we assumed that exaggeration of this response in mPges-1 mice was attributable to suppression of PGE_2_, despite their augmented formation of PGI_2_. To our surprise, deletion of the Ipr attenuated the hypertensive response to mPges-1 deletion. Furthermore, this was observed in male, but not female mice. Mechanistically, we found that Ipr deletion results in a release of the vasodilator, atrial natriuretic peptide (ANP) (15, 16) and attenuation of the oxidant stress that characterizes hyperlipidemia (17) in male mice. This results in abrogation of the hypertensive response to salt. In females, by contrast, these responses were not observed, while in ovariectomized mice estrogen attenuated salt-evoked hypertension in both Ldlr KO and Ipr/Ldlr DKO but not in mPges-1/Ldlr DKO mice.

## Results

### Deletion of the Ipr in mPges-1-deficient hyperlipidemic mice abrogates salt-evoked hypertension

Hyperlipidemic mice (Ldlr KO) were used in the current study to simulate more closely the atherosclerosis likely extant in elderly patients targeted for analgesia with mPGES-1 inhibitors. As shown in Supplemental Figures 1A-1D, despite feeding a chow diet, plasma cholesterol and/ or triglyceride levels of Ldlr KO, Ipr- and mPges-1-deficient Ldlr DKO mice were significantly elevated.

Male Ldlr KOs fed a HSD showed a time-dependent elevation of SBP in the active (night) period (Figure 1A-1B). The SBP was significantly elevated in week 2 compared with baseline during the active phase. Deletion of mPges-1 significantly increased further the salt-evoked BP response. By contrast, deletion of the Ipr unexpectedly restrained the hypertensive response to the HSD in both Ldlr KOs and mice also lacking mPges-1. At baseline, male mice lacking both mPges-1 and Ldlr had elevated BP compared to the other genotypes (Figure 1). Thereafter the attenuating effects of Ipr deletion became apparent: SBPs of Ldlr KO, mPges-1/Ldlr DKO and Ipr/mPges-1/Ldlr TKO mice were significantly elevated compared with Ipr/Ldlr DKOs one week and/ or two weeks after feeding them the HSD. Similar differences in diastolic blood pressure (DBP) responses were observed in all mutants and their littermate controls fed a HSD in the active and resting periods (Figure 1C-1D). DBP in Ldlr KOs was significantly elevated at week 2 compared with baseline during the active phase. Compared with Ipr/Ldlr DKO, the DBPs of Ldlr KO, mPges-1/Ldlr DKO and Ipr/mPges-1/Ldlr TKO mice were significantly elevated at baseline, one week and/ or two weeks after HSD feeding. However, these HSD evoked BP responses were not observed in female hyperlipidemic mice (Supplemental Figure 2A-2D). In addition, weight gain, urinary output/ fluid intake ratio and urinary sodium levels did not appear to explain the sex differences in BP responses to the salt loading in our mice (Supplemental Figure 3A-3C). We were not able to measure accurately food intake in the current study because the HSD was very hygroscopic.

**Figure 1.**
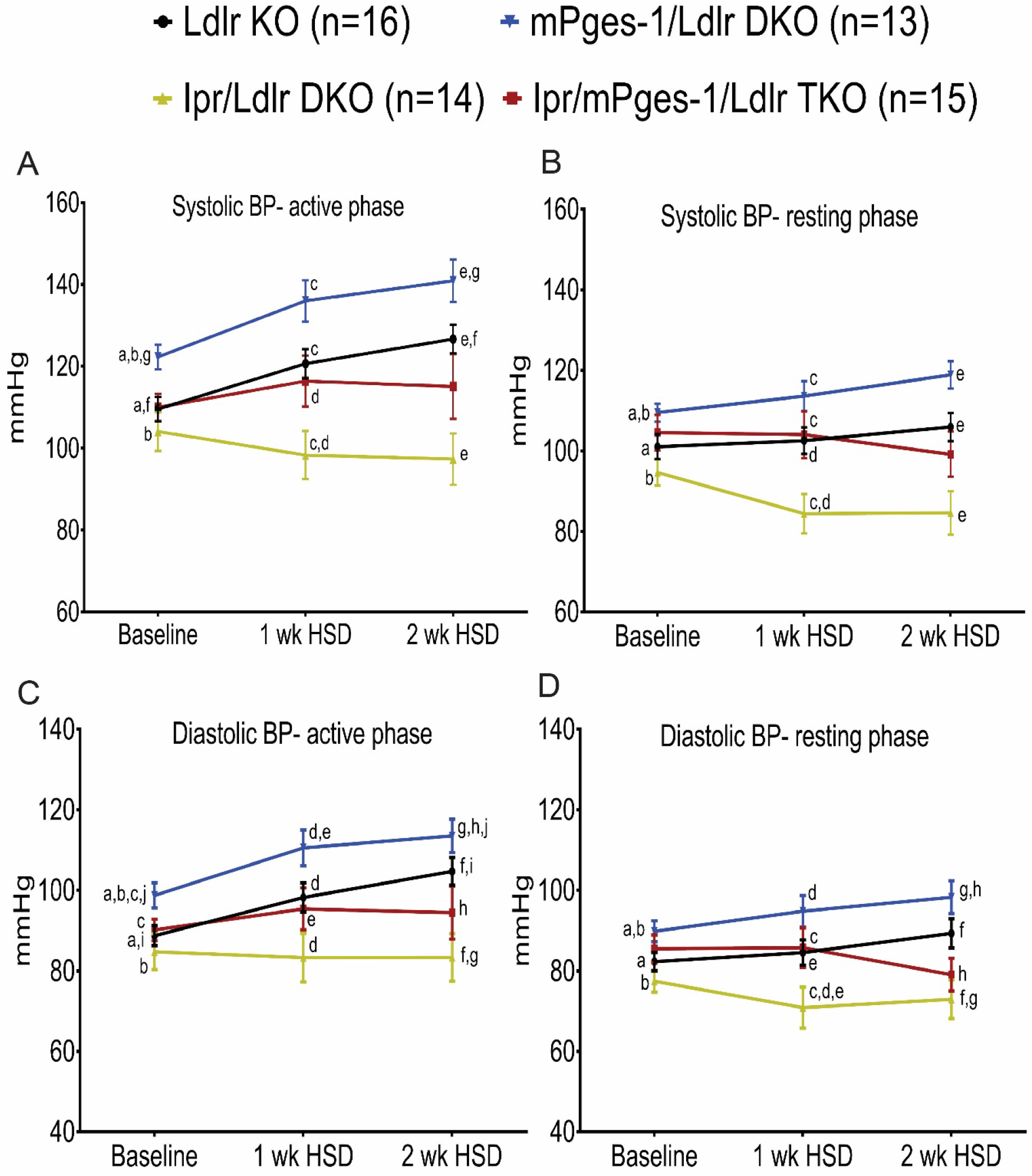
Deletion of Ipr in mPges-1-deficient male hyperlipidemic mice abrogates salt-evoked hypertension. Systolic blood pressures (SBP) in male hyperlipidemic mice and mutants fed a high salt diet (HSD) were measured via telemetry. HSD increased SBPs in Ldlr KO mice in a time-dependent pattern, during both the active (night) and resting (day) periods (A-B). Deletion of mPges-1 in Ldlr KO mice augmented salt-evoked hypertension. By contrast, deletion of prostacyclin receptor (Ipr) restrains salt-evoked hypertension and abrogated hypertensive phenotype in Ipr/mPges-1/Ldlr TKO mutants. 4-way ANOVA with repeated measures showed a significant effect of Ip, mPges-1, phase and a few of the 2- and 4-way interactions (Ipr:week, week:phase, Ipr:mPges-1:week:phase) on SBP. A posthoc pairwise t-test showed a significant effect on SBP at week 2 in respect to baseline for Ldlr KO. (C-D) Similar trends in DBP responses were observed in all mutants and their littermate controls fed an HSD both in active and resting periods. 4-way ANOVA with repeated measures showed a significant effect of Ipr, week, phase and week:phase interaction on DBP. Pairwise t-test showed significant effect on DBP only at week 2 in respect to baseline for Ldlr KO. Pairwise t-tests were used to test significant differences between Ldlr KO, Ipr/mPges-1/Ldlr TKO, Ipr/Ldlr DKO and mPges-1/Ldlr DKO. Genotypes and feeding periods with the same lower case letter are significantly different (a-j, *p*< 0.05) at baseline, 1 wk HSD or 2 wk HSD. For example, a-baseline SBP (active phase) of mPges-1/Ldlr DKO was significantly elevated compared with Ldlr KO and b-Ipr/Ldlr DKO; f-SBP (active phase) of mPges-1/Ldlr DKO was significantly elevated at 2 wk HSD compared with baseline. Data are expressed as means ± SEMs. n=13-16 per genotype.

Detailed statistical analyses of the interactions among genotypes, treatment (week) and phases for both sexes are described in supplemental Figure 4.

### Impact of Ipr and mPges-1 deletion on prostaglandin biosynthesis in male hyperlipidemic mice on a high salt diet

Two weeks of HSD feeding suppressed PGE_2_ but increased PGI_2_ biosynthesis in male Ldlr KO mice, as reflected in their urinary PGEM (7-hydroxy-5, 11-diketotetranorprostane-1, 16-dioic acid) and PGIM (2, 3-dinor 6-keto PGF_1α_) metabolites, respectively (Figure 2A-2B). Overall (Figure 2), deletion of mPges-1 in the hyperlipidemic mice (mPges-1/Ldlr DKO) suppressed PGE_2_ and augmented formation of PGI_2_, thromboxane (Tx)B_2_ and PGD_2_ as expected consequent to substrate rediversion. These changes were more pronounced on the HSD. Finally, deletion of the Ipr resulted in a reactionary increase in biosynthesis of PGI_2_, but also TxB_2_ and PGD_2_, again apparent on an HSD.

**Figure 2.**
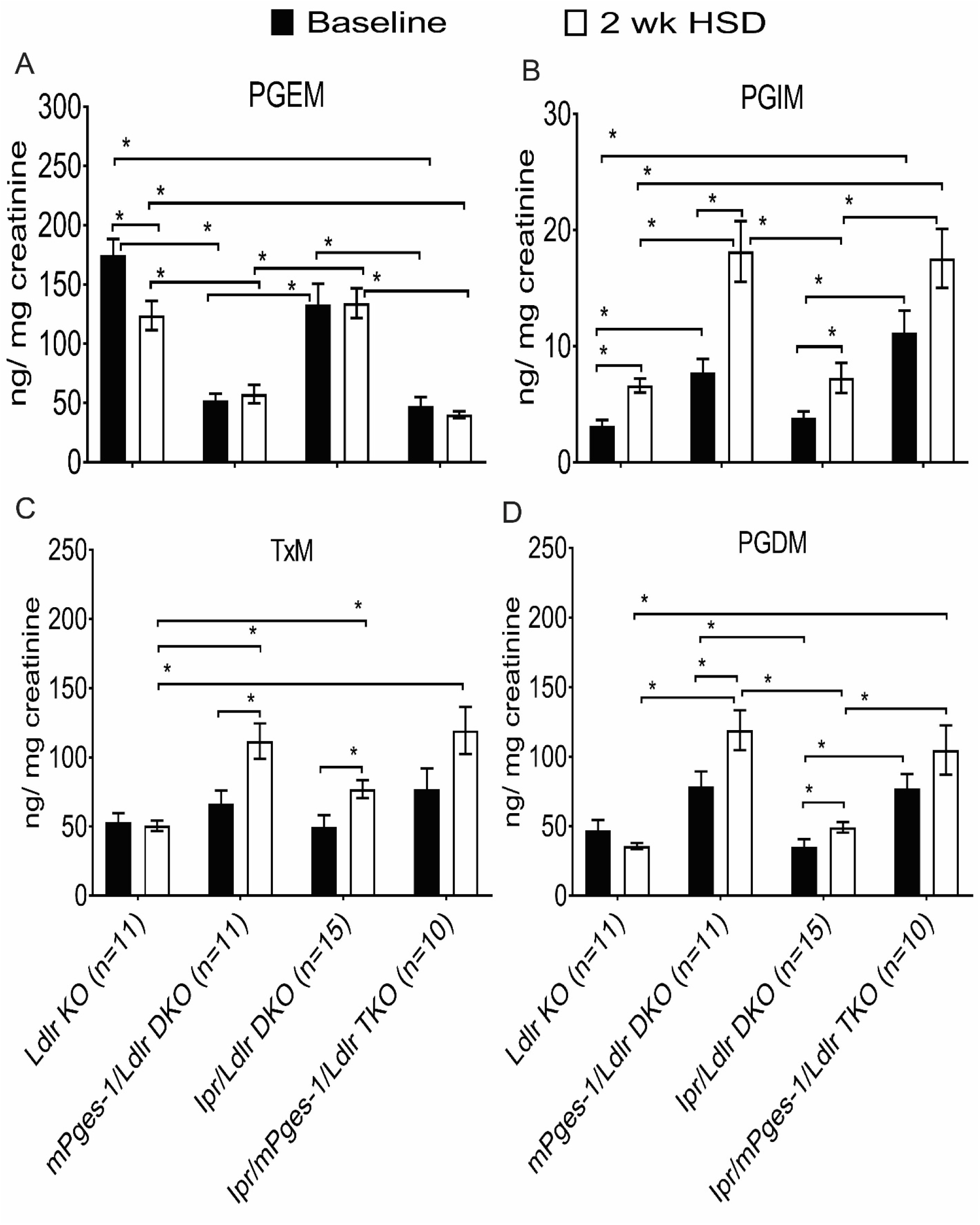
Impact of Ipr and mPges-1 deletion on prostaglandin biosynthesis in male hyperlipidemic mice on a high salt diet. Fasting (9am-4pm) urine samples from Ldlr KO, mPges-1/Ldlr DKO, Ipr/Ldlr DKO and Ipr/mPges-1/Ldlr TKO mice were collected before and two weeks after feeding a HSD and prostanoid metabolites were analyzed by liquid chromatography/ mass spectrometry, as described in the Methods. Ldlr KO mice fed an HSD suppressed PGE_2_ but increased PGI_2_ biosynthesis as reflected in their urinary PGEM (7-hydroxy-5, 11-diketotetranorprostane-1, 16-dioic acid) (A) and PGIM (2, 3-dinor 6-keto PGF_1α_) (B) metabolites, respectively. Deletion of mPges-1 suppressed PGE_2_ but increased PGI_2_ biosynthesis in mPges-1/Ldlr DKO and Ipr/mPges-1/Ldlr TKO mice. Deletion of Ipr did not alter PGEM and PGIM levels at baseline but increased PGIM on the HSD. Feeding a HSD also increased urinary 2, 3-dinor TxB_2_ (TxM) levels in DKO mutants (C). After feeding a HSD, urinary PGDM (11, 15-dioxo-9α-hydroxy-2,3,4,5-tetranorprostan-1,20-dioic acid) (D) levels were significantly elevated in the mPges-1/Ldlr DKO and Ipr/Ldlr DKOs. 3-way ANOVA showed that urinary PGIM, PGDM and TxM were significantly affected by mPges-1 deletion when mice were fed an HSD. PGEM interacted significantly alone and together with Ipr status and whether the mice were on an HSD. Pairwise t-tests were used to test for significant differences between Ldlr KO, Ipr/mPges-1/ Ldlr TKOs, Ipr/Ldlr DKO and mPges-1/Ldlr DKO. Data are expressed as means ± SEMs. \**p*< 0.05; n=10-15 per genotype.

Detailed statistical analyses of the interactions among urinary prostaglandin metabolites, genotypes and treatment (week) are described in supplemental Figure 5.

### Pharmacological inhibition of the human mPGES-1 enzyme elevates systolic blood pressure in hyperlipidemic male mice

To confirm the hypertensive phenotype of global mPges-1/Ldlr DKOs, an mPGES-1 inhibitor (MF970, 10 mg/Kg BW) was administered concomitantly with a high fat diet (HFD) for 39 weeks in humanized mPGES-1 Ldlr KO male mice. As shown in Supplemental Figure 6, inhibition of mPGES-1 suppressed urinary PGEM (A) and elevated SBP response (B) as compared to control with the HFD alone.

### A HSD activates atrial natriuretic peptide synthesis and release in Ipr-deficient mice

The unexpected suppression of the salt-evoked elevation of BP by Ipr deletion prompted us to compare gene expression profiles in the renal medullae of male Ldlr KOs and Ipr/Ldlr DKOs by high throughput RNA sequencing. We identified 897 differentially expressed genes (DEGs), with a log fold change ranging from 2.76 and −3.07 between Ldlr KO and Ipr/Ldlr DKOs at a false discovery rate cutoff of 0.033. Four hundred and forty five of these 897 DEGs were upregulated and 452 were downregulated in the renal medulla of Ipr/Ldlr DKOs compared to Ldlr KOs. Ingenuity Pathway analysis (IPA) was used to assess changes in biological pathways associated with gene expression (Table 1) and the pathways most enriched with DEGs included eukaryotic initiation factor (eIF2), eIF4/ p70S6K and mTOR signaling, mitochondrial dysfunction and oxidative phosphorylation. Thirty five of the 44 identified genes in the eIF2 pathway were downregulated in the Ipr/Ldlr DKOs, mostly members of the 60s and 40s ribosomal subunits involved in RNA binding (Figure 3A and Supplemental Table 1). Twenty three of 24 genes related to mitochondrial dysfunction and oxidative phosphorylation were downregulated in the Ipr/Ldlr DKOs (Figure 3A and Supplemental Table 1). Most of these genes are components of mitochondrial complexes I to V, which are involved in electron transport and ATP synthesis. We validated three of the genes (Downregulated; Atp5e-a subunit of mitochondrial ATP synthase. Upregulated; Cat and Sod2-antioxidant enzymes) in the mitochondrial dysfunction and oxidative phosphorylation pathways by RT-qPCR (Supplemental Figure 7A-7B). In addition, the RNA-Seq data are consistent with activation of the atrial natriuretic peptide (ANP) pathway. Expression of neprilysin (Mme) that degrades natriuretic peptides was elevated in Ipr/Ldlr DKOs compared with Ldlr KOs (Figure 3B). We confirmed by RT-qPCR analyses that mRNA levels of corin (ANP-converting enzyme) and ANP, but not brain natriuretic peptide (BNP), were significantly increased in whole heart lysate in Ipr/Ldlr DKOs (Figure 3C-3E). Moreover, renal medullary expression of Npr1, a receptor of ANP, was significantly increased (Figure 3F). Consistent with the gene expression data, urinary ANP levels were also elevated in Ipr/Ldlr DKOs compared with Ldlr KOs after two weeks on the HSD (Figure 4A-4B). We did not observe a significant difference in creatinine levels in the urine samples between Ldlr KOs and Ipr/Ldlr DKO mutants (Supplemental Figure 3D). Thus, elevated urinary ANP levels were not likely to be confounded by differences in fluid intake. Consistent with the role of PGI_2_ in restraining oxidative stress in atherosclerotic vasculature (18) and in salt-induced hypertension (19, 20) and the elevation of PGI_2_ biosynthesis on the HSD (Figure 2), excretion of a major urinary F_2_-isoprostane (F_2_iP), an index of lipid peroxidation, was not significantly elevated in Ldlr KOs after 2 weeks on a HSD (Figure 4C and 4D). However, rather than increase with Ipr deletion, F_2_iP excretion, just like BP, unexpectedly fell, consistent with the changes in mitochondrial dysfunction and oxidative phosphorylation genes observed in the renal medulla of Ipr/Ldlr DKO mice (mostly downregulated in the Ipr/Ldlr DKO) (Figure 4D). The reduction in urinary F_2_iP and elevated ANP levels consequent to Ipr deletion in the Ldlr KOs was abrogated by treatment with the ANP receptor antagonist, A71915 (21–23) (Figure 4E and 4F). This is consistent with evidence that ANP is both a vasodilator and a restraint on oxidative stress (22, 24).

**Figure 3.**
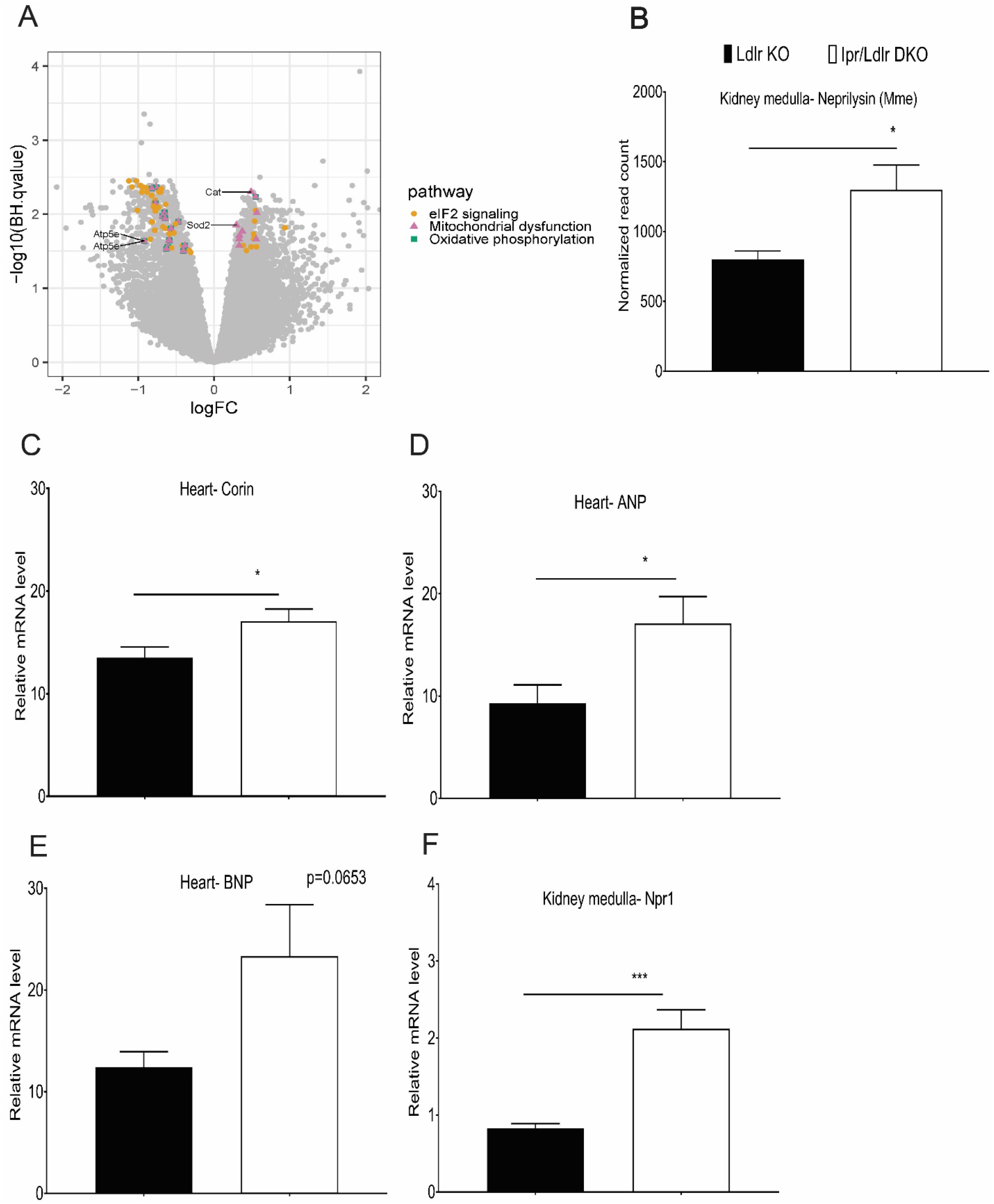
Combined Ipr deletion and salt-evoked hypertension downregulates eIF2, mitochondrial dysfunction and oxidative phosphorylation pathways and activates atrial natriuretic peptide synthesis. RNA samples isolated from kidney medulla of Ldlr KO and Ipr/Ldlr DKO after two weeks on an HSD were used for RNA-Seq. (A) Analysis of signaling pathways. A Volcano plot compares overlap of genes identified in the top three canonical pathways: eIF2 signaling, mitochondrial dysfunction and oxidative phosphorylation. 44 genes in eIF2 signaling pathway were unique. 24 genes were common between mitochondrial dysfunction and oxidative phosphorylation, and nine genes were unique to mitochondrial dysfunction. Atp5e, Cat and Sod2 are genes validated by RT-qPCR. (B) Neprilysin (Mme) transcript was increased in Ipr/Ldlr DKOs compared with Ldlr KOs. (C-F) Real-time PCR was used to measure the expression of corin (ANP-converting enzyme), atrial natriuretic peptide (ANP) and brain natriuretic peptide (BNP) in whole heart and kidney medullary Npr1 (a receptor of ANP). Feeding HSD increased corin and ANP transcripts in heart and Npr1 in kidney medulla of Ipr/Ldlr DKOs compared with Ldlr KOs. BNP gene was not significantly altered between Ldlr KO and Ipr/Ldlr DKOs. Data are expressed as means ± SEMs (Parametric test, two-tailed, \**p*< 0.05, \*\*\**p*< 0.001; n=9-10 per genotype).

**Figure 4.**
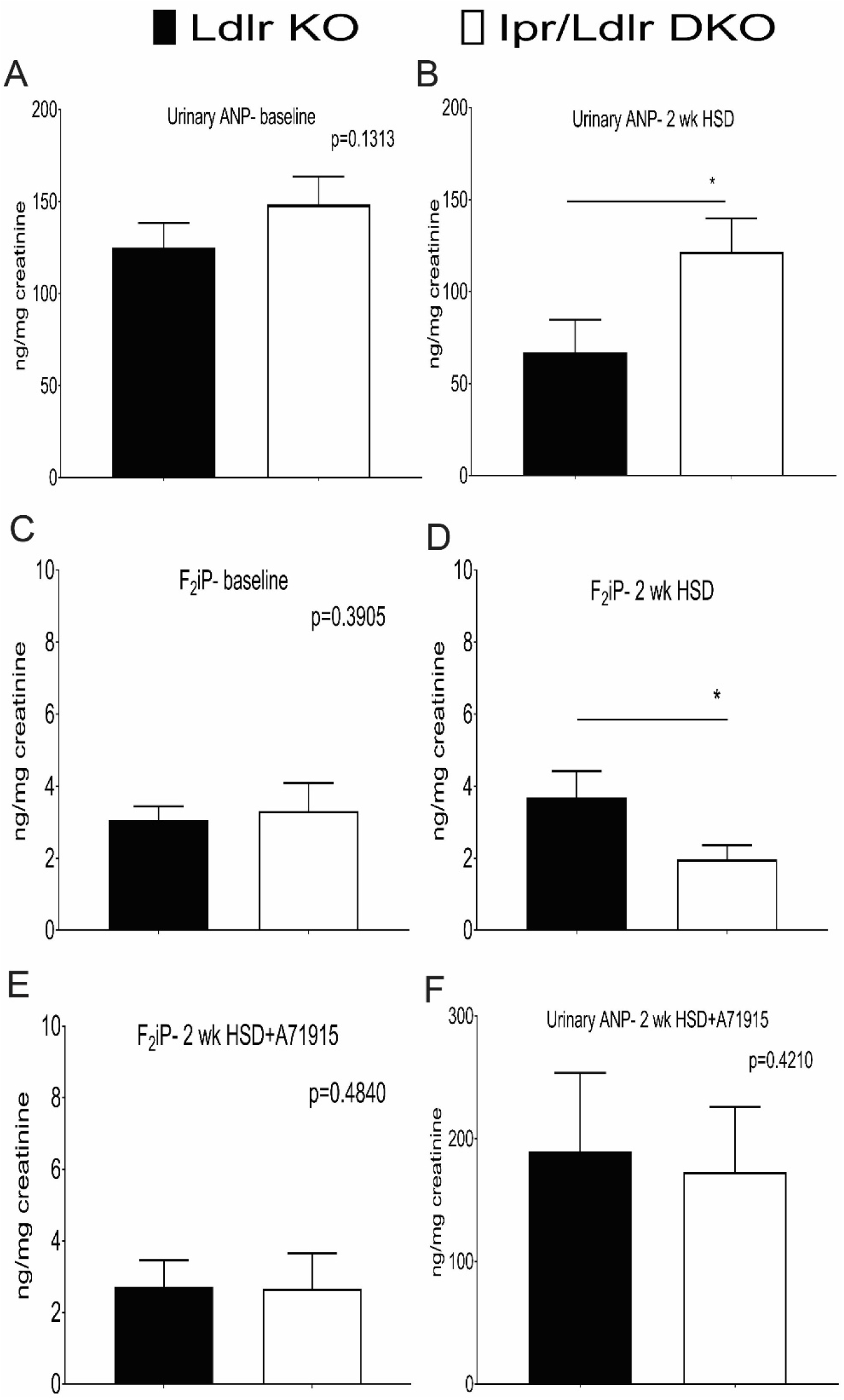
Combined deletion of Ipr and salt-evoked hypertension increases urinary atrial natriuretic peptide and reduces F_2_-isoprostanes. Urinary ANP levels at baseline (A) and two weeks after a HSD (B) were measured using an ELISA kit. Urinary ANP levels were elevated in Ipr/Ldlr DKO compared with Ldlr KOs two weeks after feeding an HSD. An abundant urinary F_2_-isoprostane (8,12-*iso*-iPF2α-VI) was analyzed by liquid chromatography/ mass spectrometry as described in the methods. (C-D) The urinary F_2_iP was not altered in Ipr/Ldlr DKO compared with Ldlr KOs at baseline, but the urinary F_2_iP in Ipr/Ldlr DKOs was significantly reduced by two weeks HSD. Treatment with ANP receptor antagonist, A71915 (50µg/Kg BW/day), abrogated the reduction of urinary ANP (E) and urinary F_2_iP (F) in Ipr/Ldlr DKO mice. Data are expressed as means ± SEMs (Parametric test, Welch’s correction, one-tailed, \**p*< 0.05; n=9-15 per genotype). We used a one-tailed test for urinary F_2_iP and ANP because both mediators had been already shown to restrain oxidative stress in the vasculature.

**Table 1.** Top Canonical Pathways Predicted by Ingenuity Pathway Analysis for the 897 Differentially Expressed Genes between Ldlr KO and Ipr/Ldlr DKO on an HSD. 57 of the 195 genes associated with eIF2 signaling pathway were differentially expressed in Ipr/Ldlr KO vs Ldlr KO.

| **Canonical Pathway** | ***P* value** | **Molecules Represented** |
| --- | --- | --- |
| eIF2 signaling | 1.87E-21 | 57/195 |
| Mitochondrial Dysfunction | 2.97E-13 | 40/157 |
| Oxidative Phosphorylation | 1.02E-11 | 29/98 |
| Regulation of eIF4 and p70S6K Signaling | 5.81E-11 | 35/146 |
| mTOR Signaling | 5.98E-08 | 35/187 |

The hypotensive phenotype of Ipr/Ldlr DKO mice was not associated with gross morphological changes in the kidney (Supplemental Figure 8) or the vasculature (Supplemental Figure 9) based on H&E staining. In male mice, deletion of the Ipr has no significant effect on urinary total nitrate + nitrite (Supplemental Figure 10A) or plasma renin levels (Supplemental Figure 10B) compared with Ldlr KO mice.

In contrast to the males, expression of corin, ANP and BNP mRNAs in whole heart and the three mitochondrial dysfunction and oxidative phosphorylation genes (Atp5e, Cat and Sod2) in the renal medulla were not significantly altered between Ldlr KOs and Ipr/Ldlr female DKOs fed the HSD for two weeks (Supplemental Figure 11A-11E). Urinary F_2_iP did not differ significantly in female Ipr/Ldlr DKOs compared to Ldlr KOs at baseline (Supplemental Figure 12A) or after two weeks on a HSD (Supplemental Figure 12B). However, combined deletion of Ipr and ANP receptor blockade in females significantly increased urinary F_2_iPs (Supplemental Figure 12C), while deletion of the Ipr significantly reduced baseline urinary ANP (Supplemental Figure 12D). This difference was abolished after two weeks on the HSD (Supplemental Figure 12E); blockade of the ANP receptor did not alter ANP levels between Ldlr KO and Ipr/Ldlr DKOs (Supplemental Figure 12F). These results were consistent with the failure of genotype to influence significantly the HSD evoked BP response in female mice (Supplemental Figure 2).

### An atrial natriuretic peptide antagonist rescues hypotension in Ipr-deficient hyperlipidemic mice on a high salt diet

Due to the physiological constraint of implanting both radio telemetry probes and mini-pumps in mice for monitoring BP and delivering the ANP antagonist during HSD feeding, we decided to use the tail-cuff system for the former while delivering the antagonist by mini-pumps. Despite being less sensitive, BP data collected with the tail-cuff system correlate with those from radio telemetry (Figure 5A-5B).

**Figure 5.**
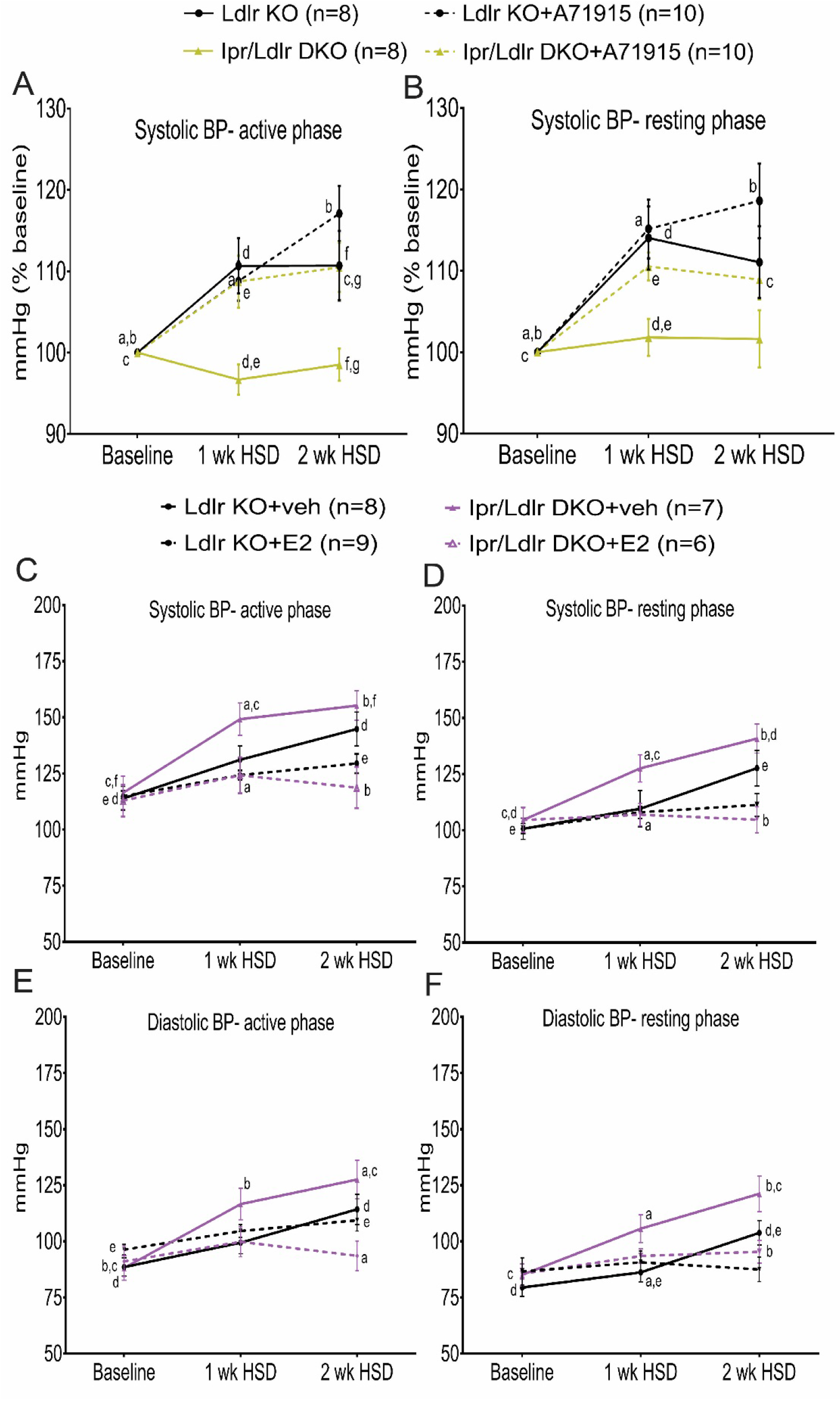
The atrial natriuretic peptide receptor antagonist (A71915) and estrogen mediate salt-evoked BP responses in Ipr-deficient male and female hyperlipidemic mice, respectively. The ANP antagonist (A71915) rescues hypotension in Ip/Ldlr DKO mice fed a HSD. Systolic blood pressures (SBP, A-active phase, B-resting phase) of male mice with and without mini-pumps were measured using a tail-cuff system before, one and two weeks after feeding a HSD in conjunction with and without ANP inhibition via A71915 infusion (50µg/Kg BW/day). To compare the effect of Ipr deletion and A71915 administration, genotype and feeding time with the same lower case letter are significantly different (a-g, *p*< 0.05) at 1 wk HSD or 2 wk HSD. For example, a-SBP (active phase) of Ipr/Ldlr DKO was significantly elevated at 1 wk HSD and b-2 wk HSD compared with baseline; d-SBP (active phase) of Ldlr KO was significantly elevated compared with Ipr/Ldlr DKO at 1 wk HSD; etc. Data are expressed as means ± SEMs. n=8-10 per group. (C-F) Salt loading increased BP in ovariectomized female Ldlr KO and Ipr/Ldlr DKO. Estradiol (E2) replacement restrained the BP responses. To compare the effect of Ipr deletion and E2 administration, genotype and/ or feeding time with the same lower case letter are significantly different (a-f, *p*< 0.05) at 1 wk HSD or 2 wk HSD. For example, a-SBP (active phase) of Ipr/Ldlr DKO+veh was significantly higher compared with Ipr/Ldlr DKO+E2 at 1 wk HSD and b-2 wk HSD; c-SBP (active phase) of Ipr/Ldlr DKO+veh was significantly higher at 1 wk HSD compared with baseline; etc. Data are expressed as means ± SEMs. n=6-9 per group.

Inhibition of the endogenous ANP signaling pathway with the antagonist, A71915 (22), attenuated the hypotensive response to Ipr deletion in the HSD fed male Ipr/Ldlr DKOs during both the active (night) and rest (day) periods (Figure 5A-5B and Supplemental Figure 13-sham-saline). Consistent with this, no significant differences in atrial and ventricular corin, ANP and BNP mRNA levels were observed between male Ldlr KOs and Ipr/Ldlr DKOs treated with the antagonist (Supplemental Figure 14A-14F). Similarly, the difference in expression of the Npr1 receptor for ANP in renal medulla (Supplemental Figure 14G), and three of the genes (Atp5e, Cat and Sod2) in the mitochondrial dysfunction and oxidative phosphorylation pathways (Supplemental Figure 15A-15C) were abolished by antagonist administration. Administration of A71915 did not alter plasma creatinine levels between male Ldlr KO and Ipr/Ldlr DKO mice (Supplemental Figure 14H). As expected in female mice, no differences in SBP or plasma creatinine were observed between Ldlr KOs and Ipr/Ldlr DKOs fed a HSD for two weeks in conjunction with ANP receptor blockade (Supplemental Figure 16A-16B).

Detailed statistical analyses of the interactions between BP, genotypes and treatment (week) in A71915 study in male mice are described in Supplemental Figure 17.

### Estrogen protects female hyperlipidemic mice from salt-evoked hypertension

To address the female BP phenotypes, we performed the HSD experiment using ovariectomized mice (OVX). HSD significantly increased BP responses in OVX Ldlr KO mice in week 2 compared with baseline during both the active and resting periods (Figure 5C-5D). Deletion of Ipr augmented the SBP responses and supplementation with estradiol (E2) significantly restrained these responses (Figure 5C-5D). Similar differences in DBP responses were observed in Ldlr KO and Ipr-deficient DKO mice (Figure 5E-5F). As expected, no significant differences in BP responses were detected among the sham-operated mice fed an HSD for 2 weeks (Supplemental Figure 19A-19D).

Detailed statistical analyses of the interactions among genotypes (Ldlr KO and Ipr/Ldlr DKO), E2 and treatment (week) of OVX mice are described in Supplemental Figure 18.

## Discussion

NSAIDs represent one of the few alternatives to opioid analgesics but themselves confer a cardiovascular hazard attributable to suppression of COX-2 derived cardioprotective prostaglandins, especially PGI_2_ (25). PGI_2_ restrains platelet activation and is a vasodilator; deletion of its Ipr receptor predisposes normolipidemic mice to thrombogenic and hypertensive stimuli (3, 26, 27). Given the importance of PGE_2_ as a mediator of pain and inflammation, interest has focused on the development of inhibitors of mPGES-1, the enzyme downstream of COX-2 that is the dominant source of PGE_2_ biosynthesis (1, 2).

In normolipidemic mice, deletion of mPges-1, unlike deletion of Cox-2 or the Ipr, has a bland adverse cardiovascular profile; it does not promote thrombogenesis and it restrains atherogenesis (3, 28). This reflects rediversion of the PGH_2_ substrate of mPGES-1 to other PG synthases, most relevantly to augment PGI_2_ biosynthesis (29). Depending on genetic background, it may leave basal and evoked BP responses unchanged or modestly increased. On a hyperlipidemic background, increased PGI_2_ limited thrombogenesis while suppression of PGE_2_ accounted for restraint of atherogenesis when mPges-1 was deleted (9). Inhibition of mPGES-1 in humans also augments biosynthesis of PGI_2_ coincident with suppression of PGE_2_ (30).

Initially, we wished to examine the impact of mPges-1 deletion on BP in hyperlipidemic mice. Both PGE_2_ and PGI_2_ may act as vasodilators and deletion of their Epr2 and Ipr receptors predisposed normolipidemic mice to HSD induced elevations of BP (26, 27, 31). Here, we found that mPges-1 deletion predisposed hyperlipidemic male, but not female mice, to the pressor response to HSD, consistent with the role of PGE_2_ in fluid volume and BP homeostasis (32). Moreover, chronic exposure to a pharmacological inhibitor (MF970) specifically targeting human mPGES-1 resulted in elevated SBP in hyperlipidemic mice on a HFD. These observations raise the possibility that despite results in healthy volunteers (28), inhibition of mPGES-1 in male patients with hyperlipidemia may predispose them to an exaggerated BP response to a HSD.

To investigate whether the augmented PGI_2_ biosynthesis resulting from mPges-1 deletion might be buffering the hypertensive phenotype we utilized mice lacking the Ipr. We were surprised to find that BP responses to salt loading in male, but not female mice, was attenuated (rather than exacerbated) in Ipr/mPges-1 DKOs. Deletion of the Ipr resulted in a compensatory increase in biosynthesis of PGI_2_ consequent to salt loading. However, given the absence of its receptor, this would be unlikely to influence directly BP homeostasis. Rather, we found activation of another compensatory mechanism, increased formation of the vasodilator ANP. Antagonism of its Npr1 receptor was sufficient to rescue the hypotensive response to a HSD in Ipr depleted mice.

Hyperlipidemia in Ldlr KOs is associated with oxidative stress, reflected by increased generation of F_2_iPs, biomarkers of lipid peroxidation (17). Both PGI_2_ and ANP can act to restrain oxidative stress which itself may contribute to elevation of BP in response to a HSD (18, 33, 34). Here we found that despite augmented biosynthesis of PGI_2_, urinary F_2_iP was depressed in Ipr/Ldlr DKOs compared to mice lacking the Ldlr alone. To address the possibility that this reflected the compensatory augmentation of ANP, we treated the mice with an ANP receptor antagonist and found that like the hypotensive phenotype, it rescued the suppression of F_2_iP in the Ipr/Ldlr DKOs. Pathway enrichment analyses of RNA-sequencing data also reflected a shift in redox balance in the renal medulla of the Ipr/Ldlr DKO mice. Some 23 genes related to mitochondrial dysfunction and oxidative phosphorylation are downregulated while genes encoding antioxidant enzymes, including mitochondrial superoxide dismutase (SOD2) and catalase, are upregulated. Again, ANP antagonism rescued this signature, adding evidence consistent with an antioxidant effect of functional relevance. Although the ANP/ Npr1 pathway plays an important role in regulating blood volume and pressure (35, 36), we failed to observe comparative diuresis or natriuresis in the Ipr/Ldlr DKO mice. Similarly, urinary total nitrate/ nitrite and plasma renin levels were unaltered in the Ipr/Ldlr DKO mice compared to Ldlr KO controls.

These differences in the BP response to a HSD and the attendant changes in gene expression and activation of the ANP pathway were observed only in male mice. There is prior evidence for the influence of sex and genetic background on disruption of the prostaglandin pathways. For example, we have shown that deletion of the Ipr accelerates atherogenesis particularly in female mice due to the importance of PGI_2_ as a mediator of estrogen receptor dependent cardioprotection (18).

Estrogen increases vasodilation partly by binding to its receptors in vascular endothelial and smooth muscle cells (SMC) of the vasculature (37). Consistent with the findings that estradiol activates PGI_2_ biosynthesis in rat aortas (38), rat aortic SMC (39) and human endothelial cells (40), ovariectomy augmented the hypertensive response to a HSD in of Ldlr KO and Ipr-deficient Ldlr DKOs. Estradiol replacement restrained the elevation in BP in the female ovariectomized mice consistent with our observation of sexual dimorphism in the response to Ipr deletion and the BP response to a HSD.

In summary, we report distinct sex dependent compensatory mechanisms to preserve BP homeostasis in response to disruption of the receptor for the direct vasodilator, PGI_2_ (Figure 6). In males, deletion of the Ipr restrains salt-evoked hypertension via activation of the ANP/Npr1 pathway reducing the oxidative stress characteristic of hyperlipidemia. In female mice, estrogen restrains the BP responses of both Ldlr KO and Ipr/Ldlr DKO mice to salt-evoked hypertension. Additionally, our findings with mPges-1 deletion or pharmacological inhibition of the enzyme in mice suggest that hyperlipidemic male patients, consuming a high salt diet, will be susceptible to hypertension when taking mPGES-1 inhibitors.

**Figure 6.**
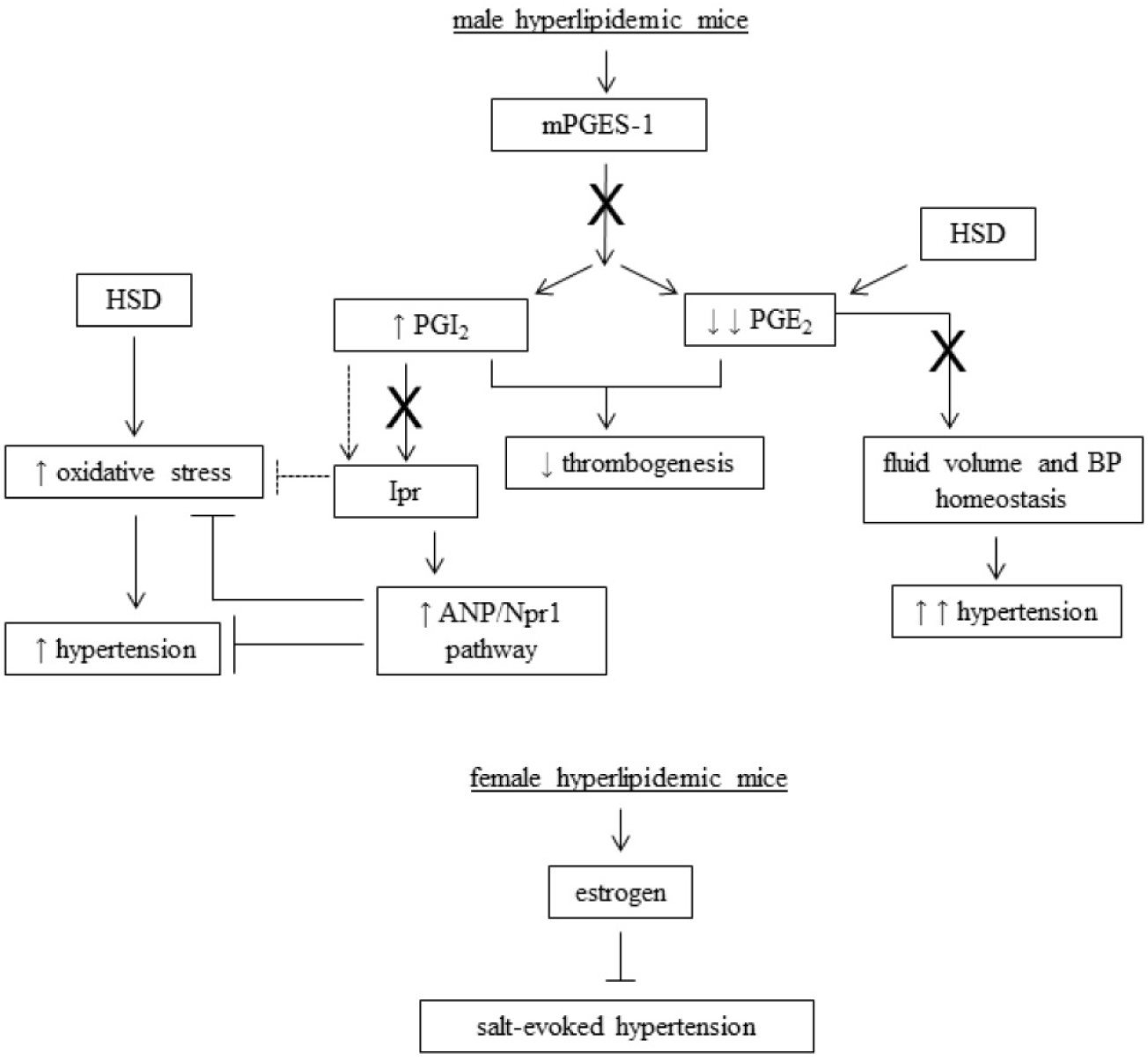
Schema depicting the effect of a high salt diet in prostaglandin I_2_ receptor deficient male and female mice. Deletion of mPges-1 suppresses PGE_2_ biosynthesis while increasing PGI_2_, which contributes to the attenuation of thrombogenesis in hyperlipidemic mice. However, salt loading suppresses PGE_2_ biosynthesis and increases BP responses. Both PGI_2_ and ANP are vasodilators and restrain oxidative stress induced by a HSD. Deletion of Ipr resulted in the compensatory increase in ANP/ Npr1 signaling and reduced mitochondrial oxidative stress and BP responses. In female hyperlipidemic mice, estrogen protects Ldlr KO and Ipr/Ldlr DKOs from salt-evoked hypertension. mPGES-1 indicates microsomal prostaglandin E synthase 1; PGE_2_, prostaglandin E_2_; PGI_2_, prostacyclin, Ipr, I prostanoid receptor; ANP-atrial natriuretic peptide; Npr1, receptor of ANP; BP, blood pressure; X, inhibition or deletion of enzyme; ̶ ̶̶ ̶̶̶̶˧, restrain; HSD, high salt diet.

## Methods

All reagents used were purchased from Sigma-Aldrich (St. Louis, MO) unless otherwise stated.

### Generation of prostacyclin receptor knockout (Ipr KO) and microsomal prostaglandin E synthase-1 knockout (mPges-1 KO) hyperlipidemic mice

Ipr and mPges-1 KOs were generated on a hyperlipidemic background by crossing them with mice lacking the low-density lipoprotein receptor (Ldlr KOs on C57BL/6 background-Jackson Laboratory) that were inter-crossed to breed Ipr ^+/-^/mPges-1 ^+/-^/Ldlr KOs. The heterozygous mice were intercrossed to produce Ldlr KOs, Ipr/Ldlr DKOs, mPges-1/Ldlr DKOs, and Ipr/mPges-1/Ldlr TKOs. Loss of Ipr, mPges-1 and Ldlr alleles were assessed by PCR analyses. Single nucleotide polymorphism (SNP) analyses showed that our mouse models achieved at least 97% purity on the C57BL/6 background. Mice of both genders were fed a high salt diet (HSD, 8% NaCl, TD.92012, Harlan Teklad, Madison, WI) and saline containing 0.9% NaCl (Hospira, Lake Forest, IL) from 8-10 weeks of age for two weeks. Mice were weighed before and after the HSD feeding. All animals in this study were housed according to the guidelines of the Institutional Animal Care and Use Committee (IACUC) of the University of Pennsylvania. All experimental protocols were approved by IACUC (protocol #804754).

### Blood pressure measurement using radio telemetry

Continuous 24-h systolic and diastolic BPs were monitored in free running mice with the Dataquest IV systems (Data Science) as described previously.(41) Briefly, after recovering from surgery, mice were kept in a 12-hour light/ dark cycle and fed a normal chow diet (0.6% NaCl). During that time, baseline systolic and diastolic BPs, heart rate and locomotor activity of mice were measured continuously for 2-3 days. After switching to a HSD, BP was again measured continuously from day 5-7 and from day 12-14. All data are expressed as averages of 12-h day or 12-h night recordings (Supplemental Figure 20).

### Fluid intake and urinary output

One week after HSD feeding, mice were housed in metabolic cages starting from 7 am for habituation in a 12-h light/ dark cycle housing facility. At 7 pm, fluid intake and urine production were recorded every 12-h for 48 hours while HSD and saline were provided *ad libitum*. Urine samples were pooled and reported as ml/g BW/day. An enrichment toy (42) was provided in each cage during the entire period.

### Blood pressure measurement using at tail-cuff system during HSD administration with an atrial natriuretic peptide receptor antagonist (A71915)

Systolic blood pressure (SBP) was measured in conscious mice using a computerized, non-invasive tail-cuff system (Visitech Systems, Apex, NC) as described (3, 9). In male mice, BP was recorded twice each day from 7 to 9 am and 7 to 9 pm for 2-3 consecutive days after 3 days of training. In female mice, BP was recorded once each day from 7 to 9 am for 3 consecutive days after 3 days of training. Averaged SBPs were reported. Atrial natriuretic peptide receptor antagonist (A71915, 50µg/Kg BW/day, H-3048, Bachem California, Torrance, CA) was dissolved in saline and delivered via minipumps in mice during HSD feeding period. A71915 was shown to inhibit cGMP response by 3 orders of magnitude in *in vitro* studies (21) and attenuated hypertension and cardiac hypertrophy in 2K1C rats with prolonged exposure to nimesulide or NS-398 (COX-2 inhibitors) via the activation of ANP signaling pathway (22).

### Pharmacological inhibition of mPGES-1 enzyme in humanized mPges-1 hyperlipidemic mice

Male mice with knock in of human mPges-1 were crossed with Ldlr KO mice and commenced at 8 weeks of age were fed a high fat diet (Harlan Teklad, TD 88137) for 39 weeks alone and in combination with a specific inhibitor of mPGES-1 (MF970, 10 mg/ Kg BW; a kind gift of Merck). BP was measured using the tail-cuff system as mentioned. Urinary prostanoid metabolites were measured by liquid chromatography/ mass spectrometry as described. Most pharmacological inhibitors of human mPGES-1 fail to engage the murine enzyme (43)

### Mass spectrometric analysis of prostanoids and isoprostanes

Urinary prostanoid metabolites were measured by liquid chromatography / mass spectrometry as described (44). Such measurements provide a noninvasive, time integrated measurement of systemic prostanoid biosynthesis (45). Briefly, mouse urine samples were collected using metabolic cages over an 8 hour period (9am to 5pm). Systemic production of PGI_2_, PGE_2_, PGD_2_, and TxA_2_ was determined by quantifying their major urinary metabolites-2, 3-dinor 6-keto PGF_1α_ (PGIM), 7-hydroxy-5, 11-diketotetranorprostane-1, 16-dioic acid (PGEM), 11, 15-dioxo-9_α_-hydroxy-2, 3, 4, 5-tetranorprostan-1, 20-dioic acid (tetranor PGDM) and 2, 3-dinor TxB_2_ (TxM), respectively. For isoprostanes (iPs), the most abundant iPs detected in urine, 8,12-*iso*-iPF_2α_-VI was measured (46). Results were normalized with urinary creatinine.

### Mass spectrometric analysis of plasma or urinary creatinine

Quantitation of plasma or urinary creatinine was performed using ultra high pressure liquid chromatography/tandem mass spectrometry (UPLC/MS/MS) with positive electrospray ionization and multiple reaction monitoring. A stable isotope-labeled internal standard (0.1mL for plasma, 1mL for urine, 2.5μg/mL [d3]-creatinine in 3% H_2_O/acetonitrile) was added to 10μL of mouse plasma or urine. The resulting solution of urine samples was further diluted by 10 times. Samples were transferred into an autosampler vial and 20 μL was injected to the UPLC-MS/MS system. A Shimadzu Prominence UPLC system was used for chromatography. The UPLC column was a 2.1 x 50 mm with 2.5μm particles (Waters XBridge BEH HILIC). The mobile phase A was 100% acetonitrile. The mobile phase B was 5mM ammonium formate (pH = 3.98). The flow rate was 350μL/min. Separations were carried out with fixed solvent gradients (12% mobile phase B). The Thermo Finnigan TSQ Quantum Ultra tandem instrument (Thermo Fisher Scientific) equipped with a triple quadrupole analyzer was operated in positive-mode ESI and the analyzer was set in the MRM mode for the analysis of creatinine. The transition for creatinine was 114>86. Quantitation was done by peak area ratio and results were normalized to the sample volume.

### Urinary atrial natriuretic peptide, total nitrate+nitrite and sodium levels and plasma renin level

Atrial natriuretic peptide (Arbor Assays, K026-H1, Ann Arbor, MI), total nitrate+nitrite (Cayman Chemical, 780001, Ann Arbor, MI) and sodium (Biovision, K391-100, Milpitas, CA) levels in mouse urine and plasma renin levels (Abcam, ab138875, Cambridge, MA) were measured following manufacturer’s instructions.

### Preparation of mouse heart and kidney medulla for real-time PCR analysis of gene expression

Briefly, heart and kidney medulla were collected from animals perfused with ice-cold PBS dissolved in UltraPure™ DEPC-treated water to minimize degradation of RNA. The tissues were immediately stored separately in RNA*later*® solution (Ambion, Austin, TX) at 4°C. After 24h, the tissue samples were transferred to −80°C for storage until analyses. RNA was extracted using TRIzol® Reagent (Life Technologies, Grand Island, NY) and RNeasy Kit (Qiagen, Valencia, CA) following the manufacturer’s protocol. The concentration and quality of extracted RNA from different tissues were measured using NanoDrop® 1000 (Thermo Scientific, Wilmington, DE) and reverse-transcribed into cDNA using Taqman Reverse Transcription Reagents (Applied Biosystems, Foster City, CA). Quantitative real time PCR was performed using Taqman Gene Expression Assays for ANP (Mm0125747_g1), BNP (Mm01255770_g1), Npr1 (Mm00435309_m1), Atp5e (Mm00445969_m1), Catalase (Mm00437992_m1) and Sod2 (Mm00449726_m1) using an ABI Prism 7900HT real-time PCR system in a 384 well plate. Results were normalized to HPRT (Mm01545399_m1).

### RNA sequencing and analysis

Total RNA from mouse kidney medulla was isolated using Trizol and Qiagen RNeasy and RNA integrity was checked on an Agilent Technologies 2100 Bioanalyzer. RNA-seq of 16 samples (8 Ldlr KOs and 8 Ip/Ldlr DKOs) was performed using the Illumina TruSeq RNA Sample Preparation Kit and SBS Kit v3 and sequenced on the Illumina HiSeq2500 system. Samples were randomized and handled in a blinded fashion during sample preparation and sequencing. Ribosomal RNA was depleted using polyA selection as part of the standard sample preparation kit.

Raw RNA-seq reads were aligned to the mouse genome build mm9 by STAR version 2.5.1b. The dataset contained an average of 15,480,386 sense and 74,242 antisense paired-end stranded 150bp reads mapping to genes, per sample. Data were normalized and quantified at both gene and exon-intron level, using a downsampling strategy implemented in the PORT pipeline (github.com/itmat/Normalization -v0.8.4-beta). Differential expression analysis was performed with Limma voom (47) to identify the most up or down regulated genes in the Ipr/Ldlr DKO, compared to the Ldlr KO. False Discovery Rate estimates were calculated using the Benjamini-Hochberg *p*-value correction. Enrichment analysis was done using the Ingenuity Knowledge Base (www.ingenuity.com). Pathway enrichment analyses was performed on the differential expressed genes (DEGs) with FDR < 0.033. Data are deposited in Gene Expression Omnibus (NCBI) under accession number (GSE115916).

### Ovariectomy

Female mice at eight weeks old were bilaterally ovariectomized following procedures as described by Souza et al (48). For the 17β-estradiol (E2) study, mice recovered from the surgery were given E2 (150nM) in saline drinking fluid before feeding a HSD for 2 weeks. During the feeding period, the concentration of E2 was reduced to 100nM due to the increase in fluid consumption (49).

### Haemotoxylin and eosin staining of kidney and aortic roots

Kidneys (n=4) and aortic roots (n=3) were formalin fixed and paraffin embedded, and 5 µm serial sections (6-10 sections) of the tissues were cut and mounted on Superfrost Plus slides for analysis of tissue morphology by HE staining.

### Statistics

All animals were the same age and on the same LdIr KO background (C57BL/6). Where conclusions involve multiple factors, two-, three- and four-way ANOVA with repeated measures was used to investigate changes in mean scores at multiple time points and differences in mean scores under multiple conditions. The residuals are normally distributed as required by ANOVA. The degrees of freedom were corrected using Greenhouse-Geisser estimates of sphericity. ANOVA tests were repeated on multiple restricted models to investigate combinations of factors’ effects. Post-hoc analysis was performed by pairwise *t*-tests, with Bonferroni correction unless otherwise stated. A significance threshold of 0.05 was used for all tests. Significance of greater than 0.01 is indicated by double-asterisks on the graphs and significance greater than 0.001is indicated by triple-asterisks unless otherwise stated. Sample sizes were based on power analysis from estimates of the variability of the measurements and the desire to detect a minimal 10% difference in the variables assessed with α = 0.05 and the power (1-β) = 0.8.

## Author contributions

SYT, STA, HM, DM and GRG contributed to acquiring and analyzing data. SYT, ER, EH and GAF contributed to data interpretation. SYT and GAF conceived of the study and are responsible for the experimental design and manuscript preparation.

## Acknowledgements

We gratefully acknowledge the advice of Dr. Matthew Palmer (Hospital of the University of Pennsylvania) on mouse kidney morphology and technical support of Weili Yan, Helen Zhou and Wenxuan Li-Feng.

