## Supplemental tables and figures for "Sex dependent compensatory mechanisms to preserve blood pressure homeostasis in PGI_2_ receptor deficient mice"

[1]*Garret A. FitzGerald, MD

Subject code: Vascular Disease

**Supplemental Table 1. Differentially expressed genes identified in eIF2 signaling, mitochondrial dysfunction and oxidative phosphorylation pathways.**

| eIF2 | | | | | | MD & OP | | | | MD | |
|---|---|---|---|---|---|---|---|---|---|---|---|
| Gene | Log FC | Gene | Log FC | Gene | Log FC | Gene | Log FC | Gene | Log FC | Gene | Log FC |
| RPS27L | -0.9766 | RPS11 | -0.7338 | RPS7 | -0.6423 | MT-ATP6 | -0.6674 | NDUFB6 | -0.57063 | OGDH | 0.330164 |
| RPS24 | -1.1262 | RPS4x | -0.5124 | RPL22 | -0.7027 | Atp5e | -0.92357 | ATP5L | -0.62977 | CAT | 0.487318 |
| ACTA2 | 0.5324 | RPL31 | -0.8256 | KL | 0.5192 | NDUFB9 | -0.64718 | NDUFA3 | -0.59299 | MT-ND6 | -0.62631 |
| RPL35A | -0.8883 | Eif2c2 | 0.9324 | RPS12 | -1.0148 | MT-CO1 | -0.44207 | MT-CO2 | -0.92286 | PSEN1 | 0.333733 |
| PIK3CB | 0.4895 | RPL14 | -0.5899 | RPS3 | -0.5915 | MT-CYTB | -0.52514 | NDUFB4 | -0.64718 | ACO2 | 0.371576 |
| RPL26 | -0.7949 | RPL27A | -0.9874 | IGF1R | 0.5203 | COX7B | -0.76509 | NDUFB5 | -0.4051 | APP | 0.561761 |
| RPS20 | -0.6776 | EIF4G1 | 0.5477 | RPL7 | -0.312 | MT-ND3 | -0.7834 | UQCRB | -0.38672 | SOD2 | 0.29209 |
| RPL12 | -0.8207 | RPL30 | -0.6971 | EIF2S3 | 0.5576 | COX15 | 0.545401 | NDUFB2 | -0.65433 | ACO1 | 0.328076 |
| RPS13 | -0.7972 | RPS25 | -0.9386 | RPS17 | -0.7751 | ATP5J | -0.47096 | MT-ND4 | -0.53893 | LRRK2 | 0.554806 |
| RPS19 | -0.8436 | RPL39 | -0.8802 | EIF3H | -0.326 | MT-CO3 | -0.92621 | | | | |
| RPL5 | -0.7353 | RPS6 | -0.6393 | EIF4G3 | 0.3819 | Cox6c | -0.61188 | | | | |
| RPL9 | -0.8927 | RPL17 | -0.9548 | MAPK1 | 0.4263 | MT-ND1 | -0.48289 | | | | |
| RPL38 | -0.7834 | RPL11 | -0.5631 | RPS23 | -0.7784 | NDUFS4 | -0.65687 | | | | |
| Rpl22l1 | -1.0835 | RPL21 | -0.4185 | RPS27A | -1.0278 | NDUFA4 | -0.82047 | | | | |
| RPS15A | -0.8789 | Rpl36a | -0.5334 | | | COX7A2 | -0.77354 | | | | |

eIF2, eukaryotic initiation factor, MD, mitochondrial dysfunction, OP, oxidative phosphorylation

**Supplemental Figures**

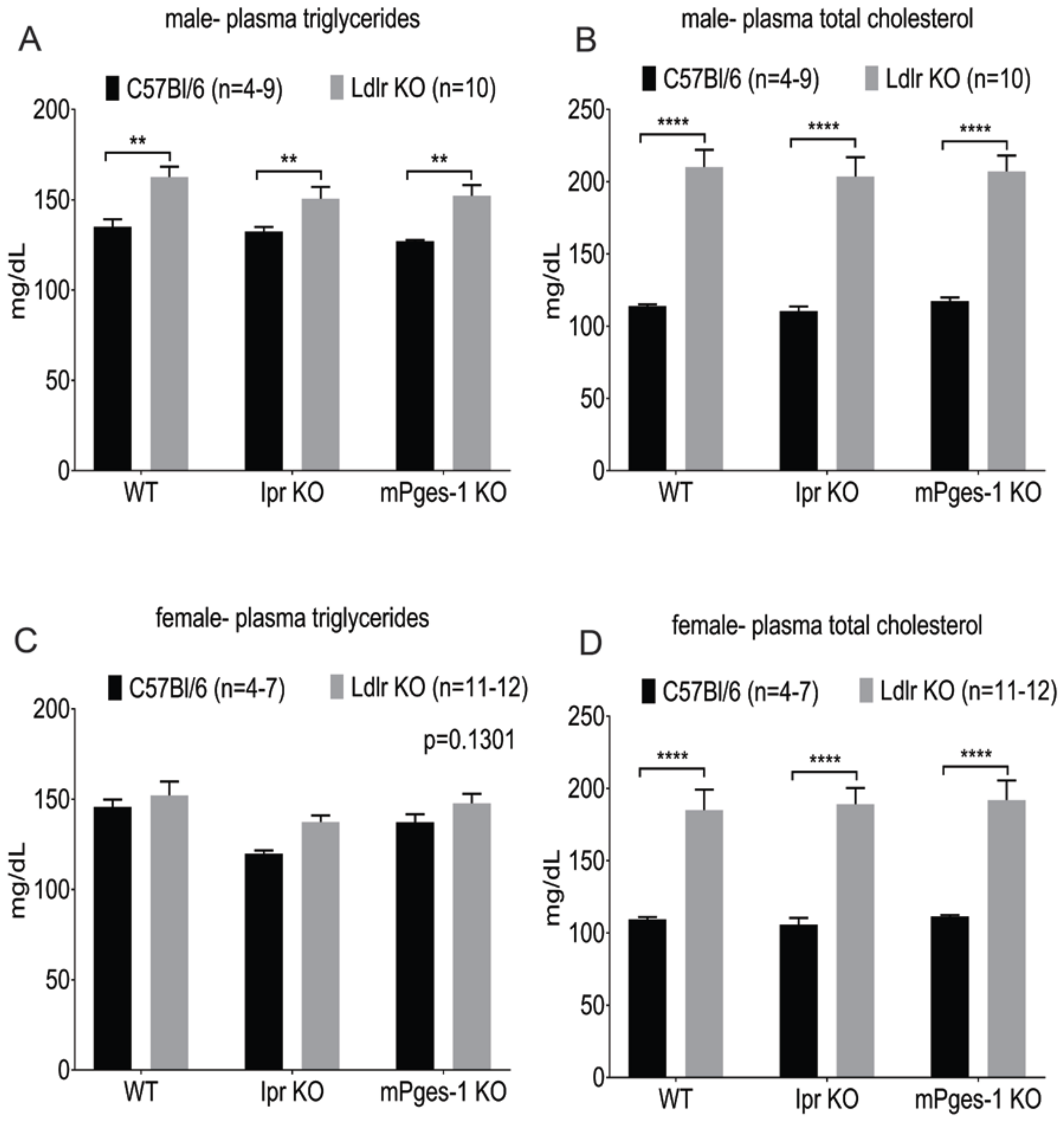


**Supplemental Figure 1. Low density lipoprotein receptor (Ldlr)-deficient mice were hyperlipidemia.** Ipr- and mPges-1-deficient mice of both sexes on C57Bl/6 and Ldlr KO genetic backgrounds between eight to ten weeks old on chow diet were sacrificed for blood collection. Plasma samples were separated and used for measurement of triglyceride and total cholesterol levels following manufacturer's instructions. A parametric t-test (2 tailed) revealed a significant effect of Ldlr KO on plasma triglycerides (A) and total cholesterol (B) levels in male wild-type (WT), Ipr- and mPges-1-deficient mice. In female mice, plasma total cholesterol levels (D) were significantly elevated in mice on the Ldlr KO background but not plasma triglyceride levels (C). Data are expressed as means ± SEMs. ***$p<0.01$, ****$p<0.0001$; n= 4-12 per group.

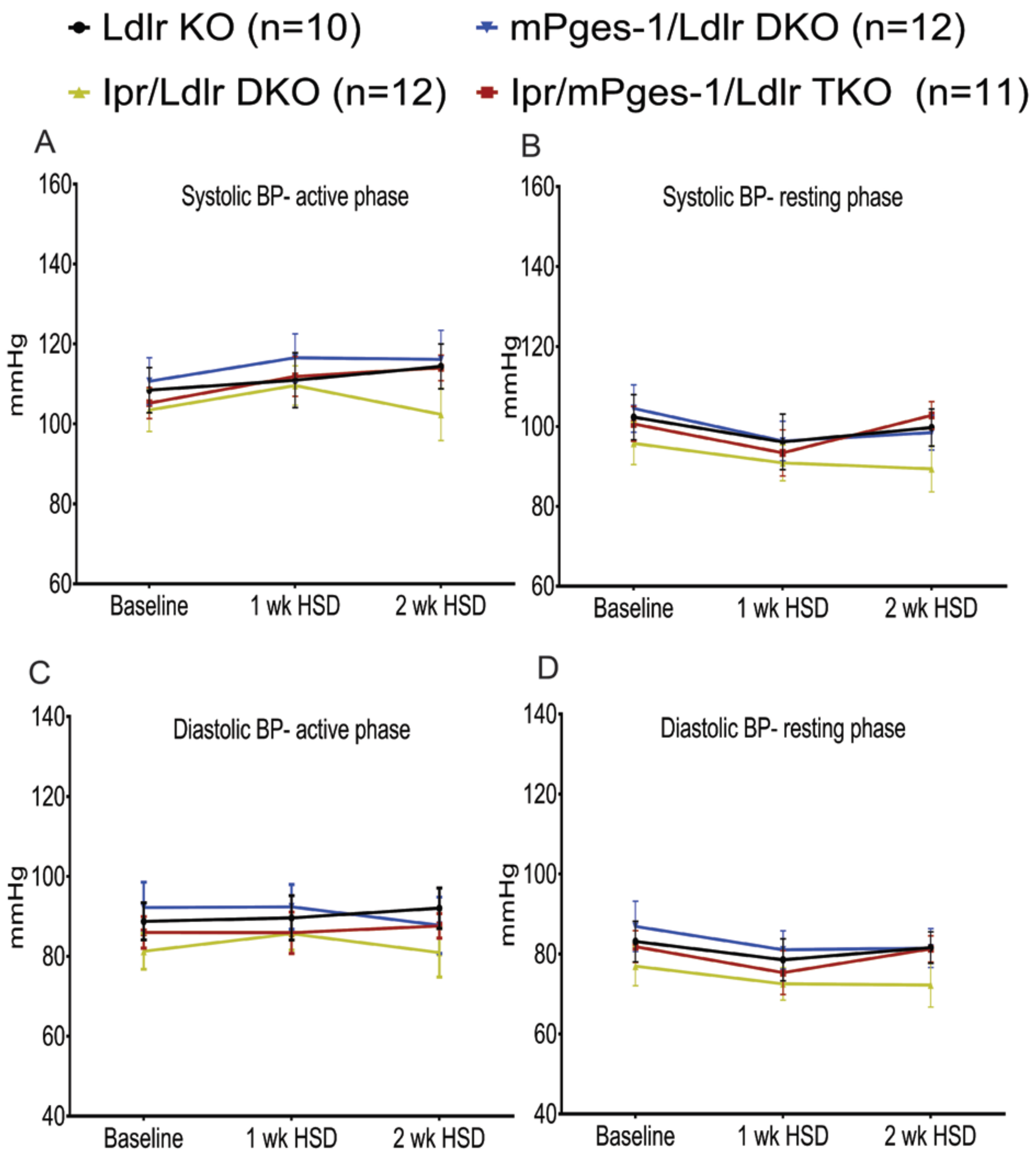


**Supplemental Figure 2. Salt loading did not alter blood pressures in female hyperlipidemic mice.** Systolic blood pressures (SBP) in female hyperlipidemic mice and Ipr-, mPges-1- and Ipr/mPges-1-deficient mutants on Ldlr KO background fed a high salt diet (HSD) were measured via telemetry. HSD did not significantly alter SBP (A and B) and DBP (C and D) in the mutants and their littermate controls (Ldlr KO), during the active phase (A and C, night) and resting phase (B and D, day). 4-way ANOVA with repeated measures and Greenhouse-Geisser correction showed no significant effects of Ipr ($p > 0.05$), treatment ($p > 0.05$), mPges-1 ($p > 0.05$) and week ($p > 0.05$) on salt-evoked elevation in SBP and DBP, although phase ($p = 0$) is significantly affecting both SBP and DBP. Data are expressed as means ± SEMs. n=10-12 per genotype.

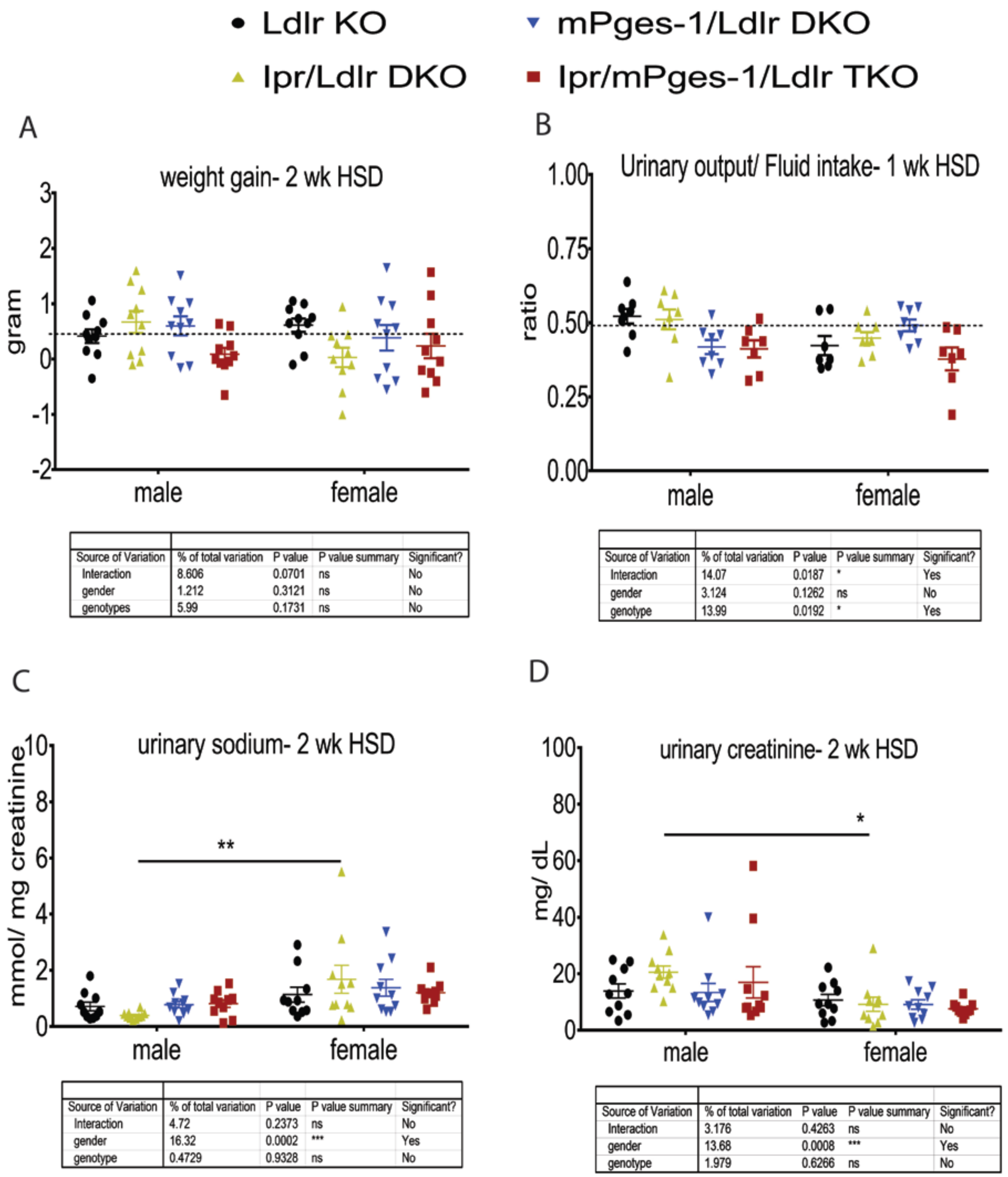


A

| Source of Variation | % of total variation | P value | P value summary | Significant? |
|---|---|---|---|---|
| Interaction | 8.606 | 0.0701 | ns | No |
| gender | 1.212 | 0.3121 | ns | No |
| genotypes | 5.99 | 0.1731 | ns | No |

B

| Source of Variation | % of total variation | P value | P value summary | Significant? |
|---|---|---|---|---|
| Interaction | 14.07 | 0.0187 | * | Yes |
| gender | 3.124 | 0.1262 | ns | No |
| genotype | 13.99 | 0.0192 | * | Yes |

C

| Source of Variation | % of total variation | P value | P value summary | Significant? |
|---|---|---|---|---|
| Interaction | 4.72 | 0.2373 | ns | No |
| gender | 16.32 | 0.0002 | *** | Yes |
| genotype | 0.4729 | 0.9328 | ns | No |

D

| Source of Variation | % of total variation | P value | P value summary | Significant? |
|---|---|---|---|---|
| Interaction | 3.176 | 0.4263 | ns | No |
| gender | 13.68 | 0.0008 | *** | Yes |
| genotype | 1.979 | 0.6266 | ns | No |

**Supplemental Figure 3. Weight gain, urinary output/ fluid ratio, urinary sodium and creatinine levels between male and female mice**. Mice were weighed before and after feeding an HSD for 2 weeks. Urinary output and fluid intake was performed at week 1 during the HSD to ensure consistency in harvesting tissues, urine and blood at the end of week 2. Urinary sodium levels were measured following manufacturer's instructions. Urinary creatinine was measured using LC-MS as described in the Methods. Two-way ANOVA revealed no significant effect of gender on weight gain (A) and urinary output/ fluid intake ratio (B). Urinary sodium (C) and creatinine levels (D) of female Ipr/Ldlr DKO mice was significantly higher and lower than male Ipr/Ldlr KOs, respectively. Multiple comparison tests (Holm-Sidak) were used to test significant differences of weight gain between male and female mice. Data are expressed as means ± SEMs. **p*<0.05, ***p*<0.01; n=7-10 per genotype.

**Supplemental Figure 4. Statistical analyses of interactions between SBP/ DBP, genotypes, treatment (week) and phases in male hyperlipidemic mice**. 4-way ANOVA with repeated measures and Greenhouse-Geisser sphericity correction showed how SBP changes for different genotypes across time at active/inactive phase, where Ipr and mPges-1 are the between-subjects factors while week and phase are the repeated measures. ANOVA indicated that SBP is significantly affected by Ipr, mPges-1, phase, and the Ipr:week and week:phase interactions. The significant factors and interactions are listed below.

- Ipr, F(1,54) = 17.2672925, p = 0.0001166
- mPges1, F(1,54) = 11.6363337, p = 0.0012316
- phase, F(1, 54) = 276.1276325, p = 0.0000000
- Ipr:week, F(2,108) = 10.0578885, p = 0.0005762
- week:phase, F(2,108) = 27.7076808, p = 0.0000000

The week factor breaks the sphericity of the variances, so we performed a 3-way ANOVA with repeated measures to detect significant differences per week. The analysis showed that Ipr and mPges-1 state, as well as phase have significant effects on SBP across all weeks, while Ipr state and phase are interacting significantly only for week 1. We also tested for differences across the different genotypes by 2-way ANOVA with repeated measures and Greenhouse-Geisser correction. This restricted model indicates that when Ipr and mPges-1 are both knocked out, the phase and the phase:week interaction affect the SBP, whereas when they are both present, the week factor is also affecting the SBP. With Ipr knocked out only, phase significantly changes SBP, while with mPges-1 knock out, both week and phase significantly affect SPB.

Similarly, 4-way ANOVA with repeated measures and Greenhouse-Geisser sphericity correction showed how DBP changes for different genotypes across time at active/inactive phase, where Ipr and mPges-1 are the between-subjects factors while week and phase are the repeated measures. ANOVA indicated that DBP is significantly affected by all four factors, and the Ipr:week and week:phase interactions. The significant factors and interactions are listed below.

- Ipr, $F(1,54) = 13.3603111$, $p = 0.0005829$
- mPges1, $F(1,54) = 7.8658849$, $p = 0.0069871$
- week, $F(2,108) = 4.1848207$, $p = 0.0241011$
- phase, $F(1, 54) = 121.9086652$, $p = 0.0000000$
- Ipr:week, $F(2,108) = 8.1299886$, $p = 0.0011381$
- week:phase, $F(2,108) = 12.5879864$, $p = 0.0000181$

A 3-way ANOVA with repeated measures was performed to detect the significant differences per week, and showed that Ipr and phase have significant effects on DBP across all weeks, with Ipr state and phase interacting significantly only at week 1. We also tested for differences across the different genotypes, by 2-way ANOVA with repeated measures and Greenhouse-Geisser correction. This restricted model indicates that phase and week:phase interaction affect the DBP in all genotypes, and when Ipr and mPges-1 are both present, the week factor has a significant effect on DBP as well. 2-way ANOVA with repeated measures for each pair of mPges-1 state and week indicates that when mPges-1 is knocked out, phase has a significant effect on DBP for all weeks. While when mPges-1 is present, the interaction of Ipr:phase is significant as well. This is also shown by 2-way ANOVA with repeated measures for each pair of mPges-1 state and phase that indicate that when mPges-1 is knocked out, a significant effect of Ipr and week is noticed during active phase only. When mPges-1 is present, Ipr state affects the DBP during resting phase, but during active phase DBP is affected by Ipr and Ipr:week interaction.

**Statistical analyses of interactions among SBP/DBP, genotypes, treatment (week) and phases in female hyperlipidemic mice**. 4-way ANOVA with repeated measures and Greenhouse-Geisser correction indicated that both SBP and DBP are significantly affected by phase ($p=0$) and the interaction week:phase ($p=0$). A 3-way ANOVA with repeated measures was performed to detect the significant differences per week, and showed that phase has significant effects on SBP and DBP across all weeks. We also tested for differences across the different genotypes, by 2-way ANOVA with repeated measures and Greenhouse-Geisser correction. This restricted model indicates that phase affects the SBP and DBP in all genotypes, and the week:phase interaction has a significant effect on DBP when Ipr is knocked out, and on SBP when either one or both of Ipr and mPges-1 are knocked out.

**Supplemental Figure 5. Statistical analyses of interactions among urinary prostaglandin metabolites, treatment (week) and genotypes**. The 3-way ANOVA with repeated measures showed how prostaglandin biosynthesis changes for different genotypes with different treatments, and specifically that PGIM, PGDM and TxM excretion is significantly affected by mPges-1 presence or absence, whether the mice are on HSD or not and the interaction of these two factors, as shown below. PGEM biosynthesis is also affected by the interaction of the Ipr state and whether the mice are on HSD or not, as well as by the 3way interaction of mPges-1 state, Ipr state and whether the mice are on HSD or not.

For PGDM: mPges1, $F(1,43)=77.4094975$, $p=0.00000000003620496$, treatment, $F(1,43)=5.5950815$, $p=0.02259109$, mPges1:treatment, $F(1,43)=4.6867501$, $p=0.03599118$

For PGEM: mPges1, $F(1,43)=67.30982657$, $p=0.0000000002439751$, treatment, $F(1,43)=5.02527221$, $p=0.03019348$, Ipr:treatment, $F(1,43)=4.69330814$, $p=0.03586802$, mPges1:treatment, $F(1,43)=4.36810887$, $p=0.04256714$, Ipr:mPges1:treatment, $F(1,43)=9.25581947$, $p=0.003991717$

For PGIM: mPges1, $F(1,43)=60.09688$, $p=0.000000001066618$, treatment, $F(1,43)=29.3896179$, $p=0.000002522259$, mPges1:treatment, $F(1,43)=5.2963732$, $p=0.02627928$

For TxM: mPges1, $F(1,43)=16.21830969$, $p=0.0002254031$, treatment, $F(1,43)=29.70607333$, $p=0.000002289881$, mPges1:treatment, $F(1,43)=8.96320747$, $p=0.004553328$

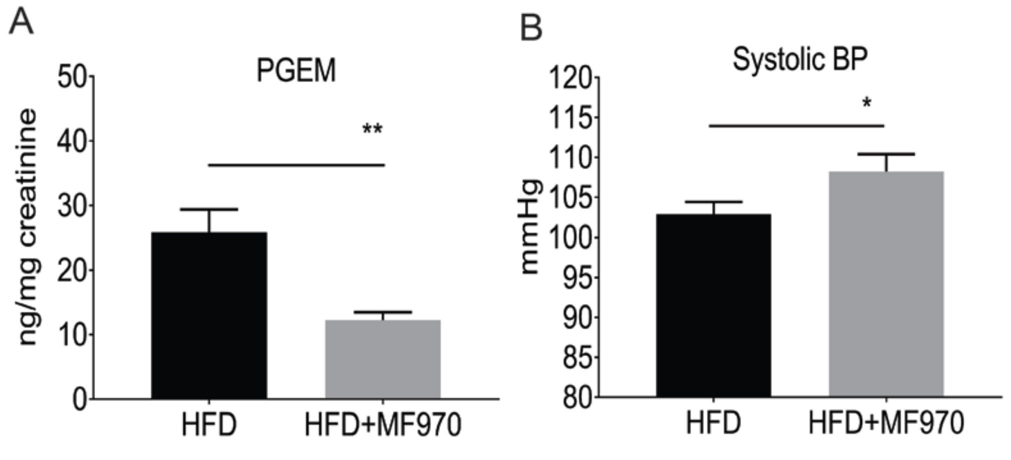


**Supplemental Figure 6. Pharmacological inhibition of mPGES-1 enzyme suppresses $PGE_2$ biosynthesis and elevates systolic blood pressure.** Eight weeks old male humanized mPges-1

hyperlipidemic mice were put on a high fat diet (HFD) alone or in conjunction with an inhibitor specific for human mPGES-1 enzyme (MF970) for 39 weeks. At the end of the HFD, blood pressure was measured using a tail-cuff system and urine samples were collected for prostanoid metabolites measurement by LC/ MS as detailed in the Methods. MF970 administration suppressed urinary $PGE_2$ metabolite (A) and elevated SBP (B). Data are expressed as means ± SEMs (Parametric test, one-tailed, *$p < 0.05$, **$p < 0.01$). n=8-15 per group. We used a one-tailed test for urinary PGEM and systolic BP because MF970 has been shown to suppress $PGE_2$ biosynthesis and $PGE_2$ regulates BP homeostasis.

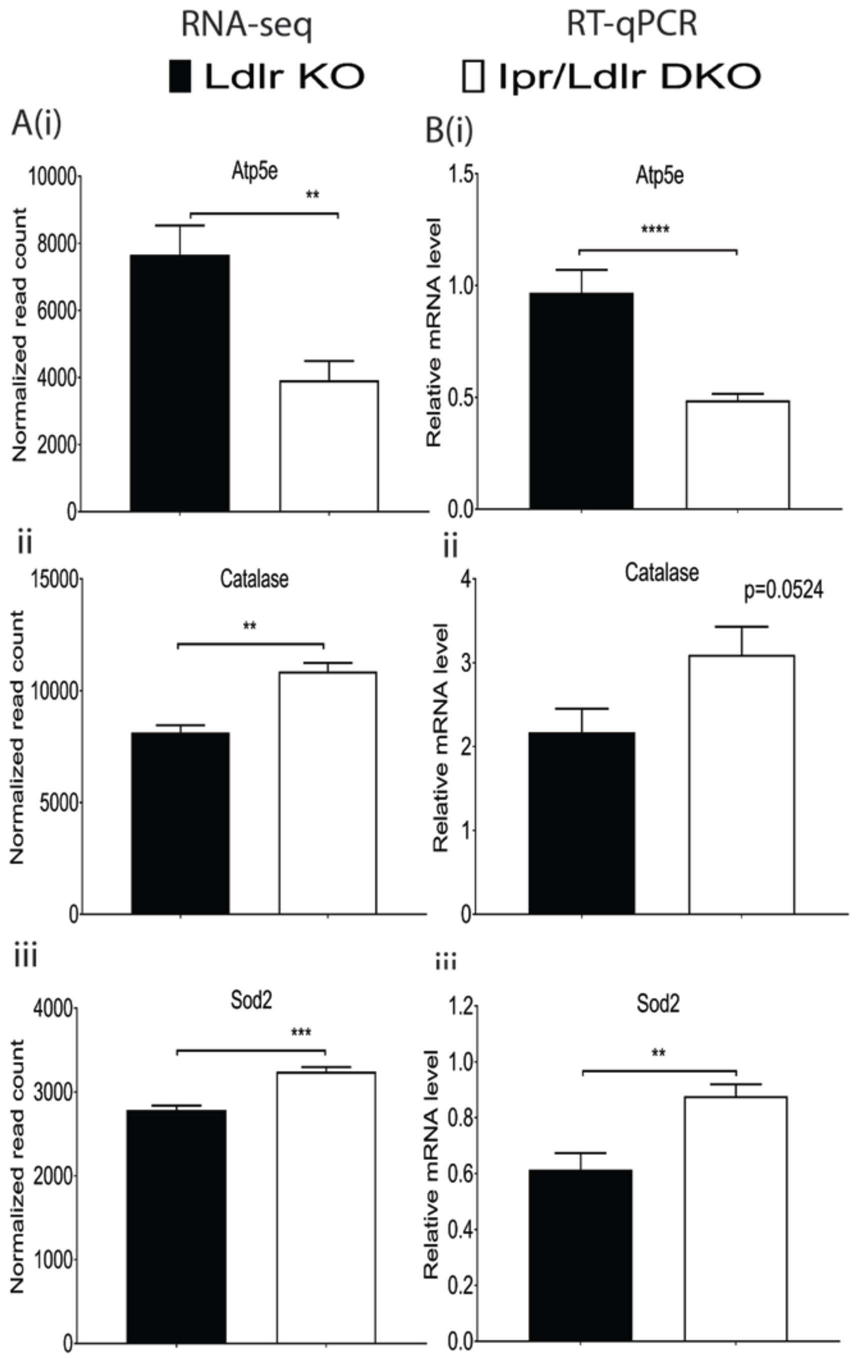


**Supplemental Figure 7. Validation of RNA-Seq transcripts in kidney medulla of male mice by RT-qPCR.** RNA samples isolated from kidney medulla of Ldlr KO and Ipr/Ldlr DKO after two weeks on an HSD were used for real-time PCR analyses. RNA-seq transcripts of three genes (A- Atp5e, catalase and Sod2) in mitochondrial dysfunction and oxidative phosphorylation pathways were consistent with their mRNA levels as detected by RT-qPCR (B). Data are expressed as means ± SEMs (Parametric test, two-tailed, **$p$<0.01, ***$p$<0.001, ****$p$<0.0001). n=8 per genotype for RNA-seq, n=10 per genotype for RT-qPCR.

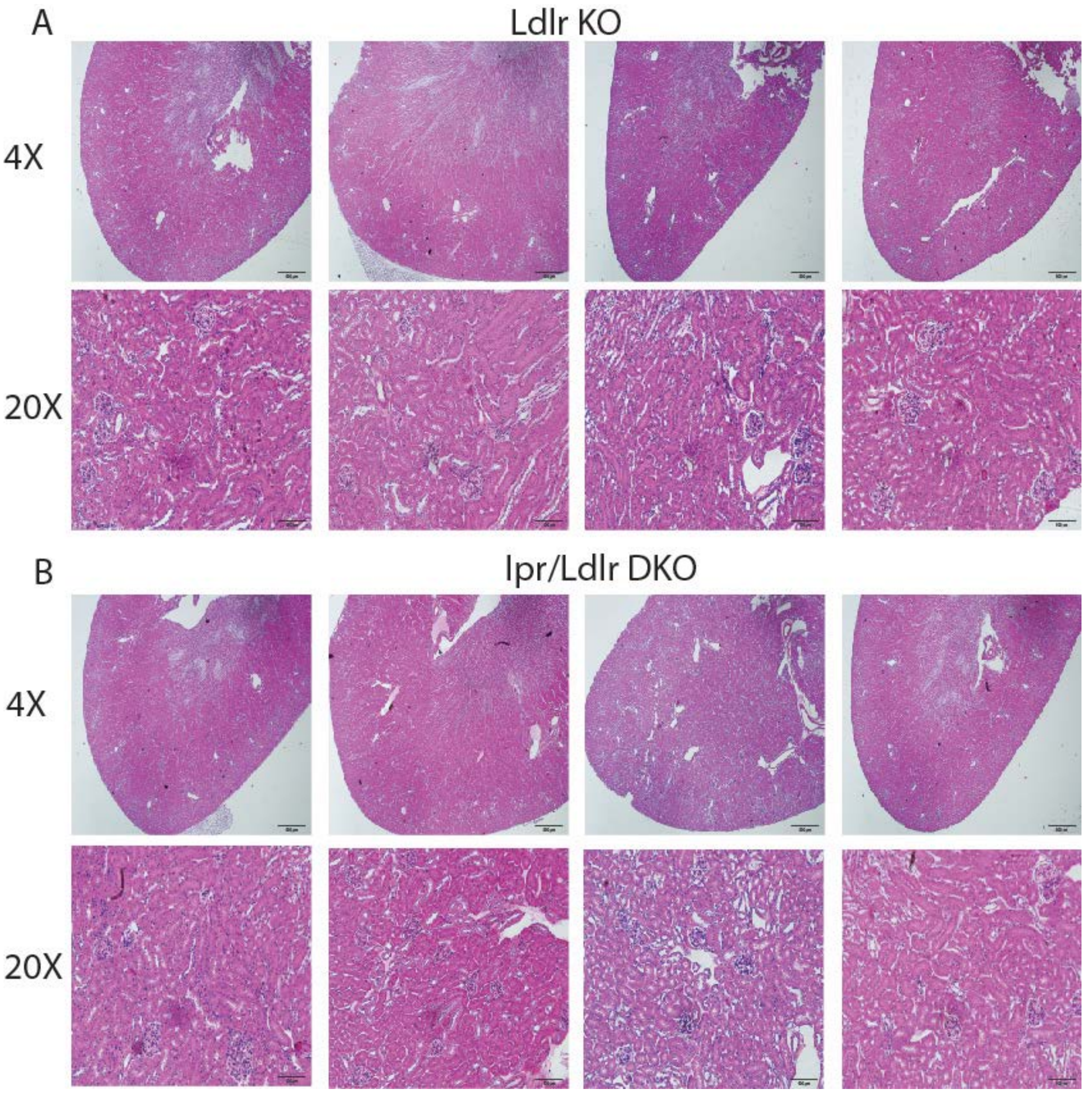


**Supplemental Figure 8. Morphological characteristics of kidney of male Ldlr KO and Ipr/Ldlr DKO mice fed a high salt diet for two weeks.**

After feeding an HSD for 2 weeks, kidneys of male Ldlr KO (A) and Ipr/Ldlr DKO (B) mice were formalin fixed and paraffin embedded. Five µm serial sections (6-8 sections) of the tissues were cut and mounted on Superfrost Plus slides for analysis of tissue morphology by HE staining. n= 4 per genotype.

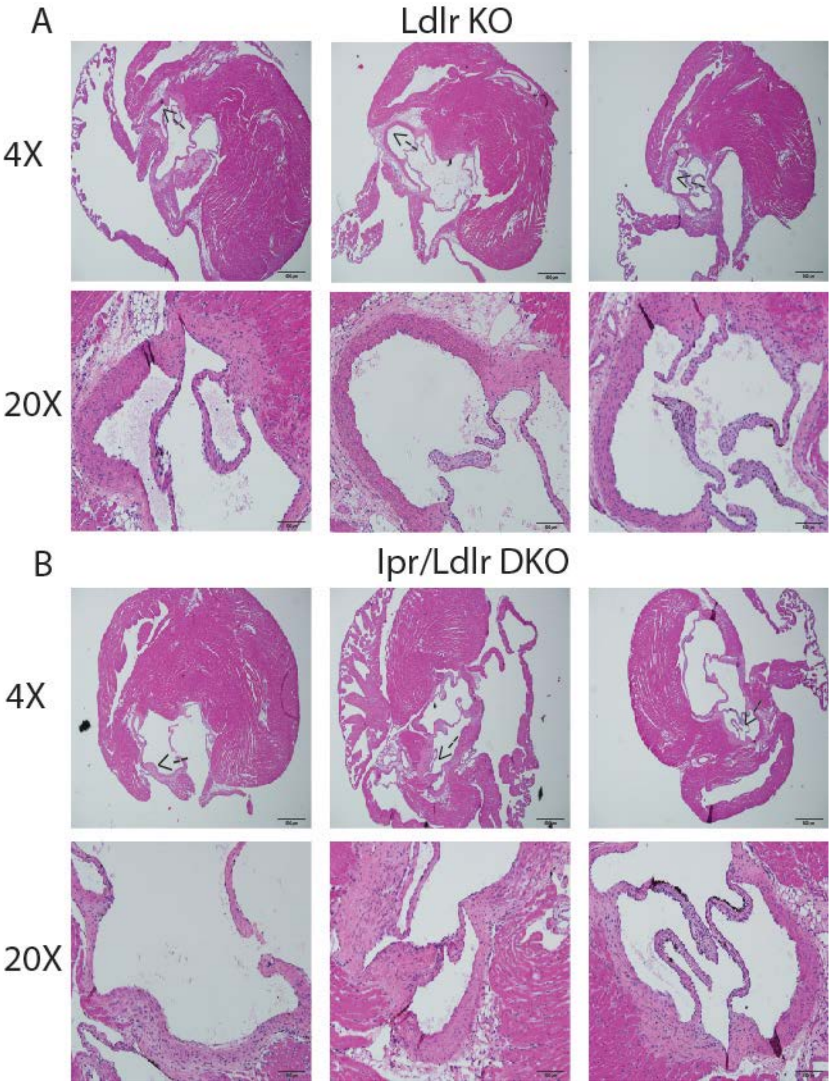


**Supplemental Figure 9. Morphological characteristics of vasculature of male Ldlr KO and Ipr/Ldlr DKO mice fed a high salt diet for two weeks.**

After feeding an HSD for 2 weeks, aortic roots of male Ldlr KO (A) and Ipr/Ldlr DKO (B) mice were formalin fixed and paraffin embedded. Five µm serial sections (8-10 sections) of the tissues were cut and mounted on Superfrost Plus slides for analysis of tissue morphology by HE staining. n= 3 per genotype. → indicates area of 20X magnification.

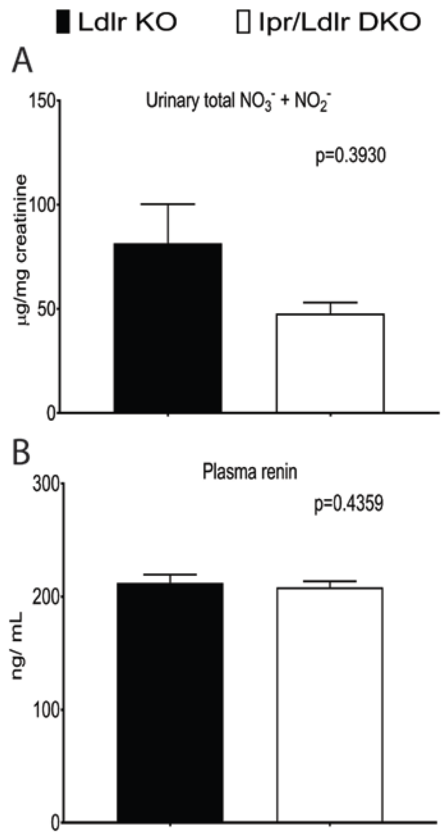


**Supplemental Figure 10. Impact of Ipr deletion on urinary total nitrate+ nitrite and sodium and plasma renin levels in male hyperlipidemic mice on a high salt diet.** Fasting (9am-4pm) urine samples from Ldlr KO and Ipr/Ldlr DKO mice were collected two weeks after feeding an HSD. Deletion of Ipr in Ldlr KO mice did not significantly alter urinary total nitrate+ nitrite (A) and plasma renin (B). Data are expressed as means ± SEMs (Parametric test, two-tailed, $p > 0.05$, n=10 per genotype).

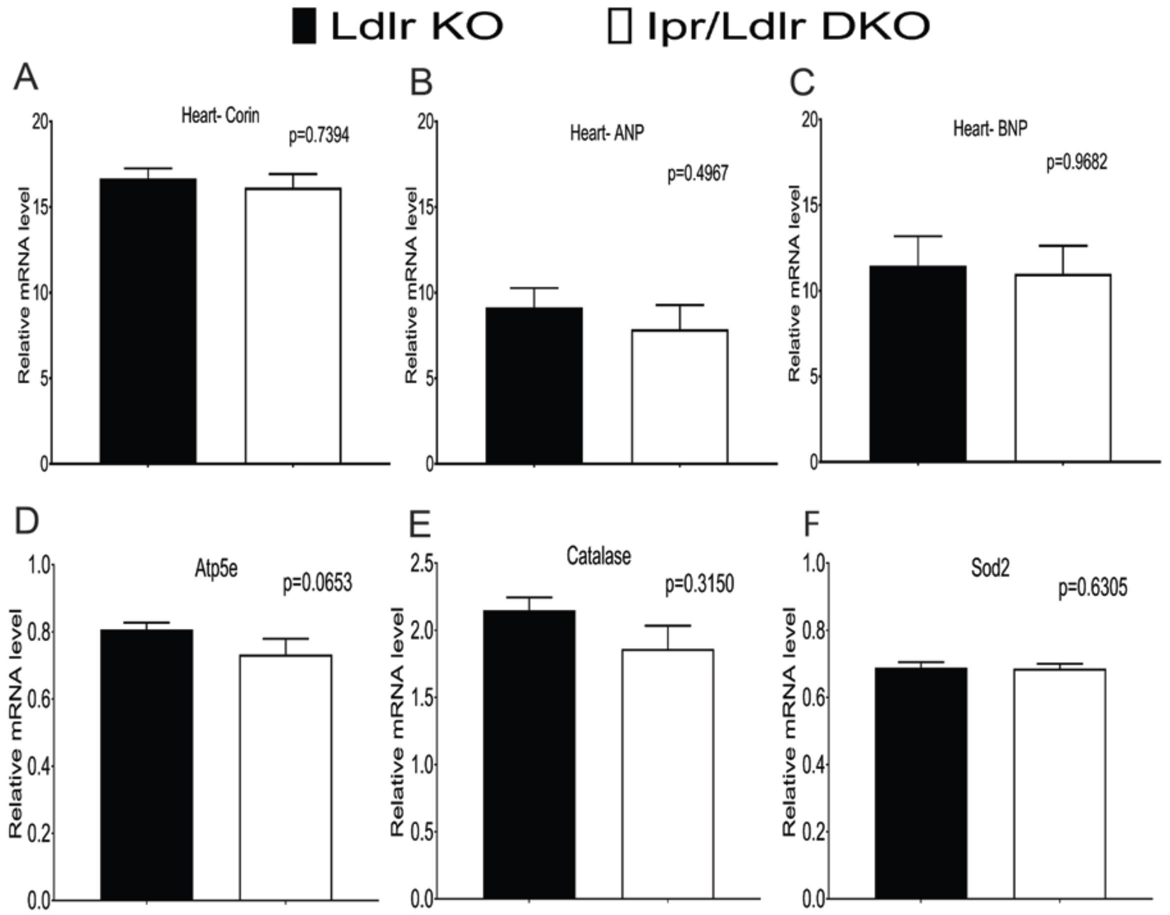


**Supplemental Figure 11. Deletion of Ipr did not alter mRNA expression of corin, atrial natriuretic peptide and brain natriuretic peptide in heart and ATP5e, catalase and superoxide dismutase 2 in kidney medulla of female mice fed a high salt diet.** RNA samples isolated from whole heart and kidney medulla of Ldlr KO and Ipr/Ldlr DKO after two weeks on an HSD were used for real-time PCR analyses. No significant differences in corin (A- ANP converting enzyme), atrial natriuretic peptide (B- heart ANP), brain natriuretic peptide (C- heart BNP) and kidney medullary Atp5e (D), Cat (E) and Sod2 (F) were observed between Ldlr KO and Ipr/ldlr DKO mice. Data are expressed as means ± SEMs (Parametric test, two-tailed, $p >$ 0.05; n=9-10 per genotype).

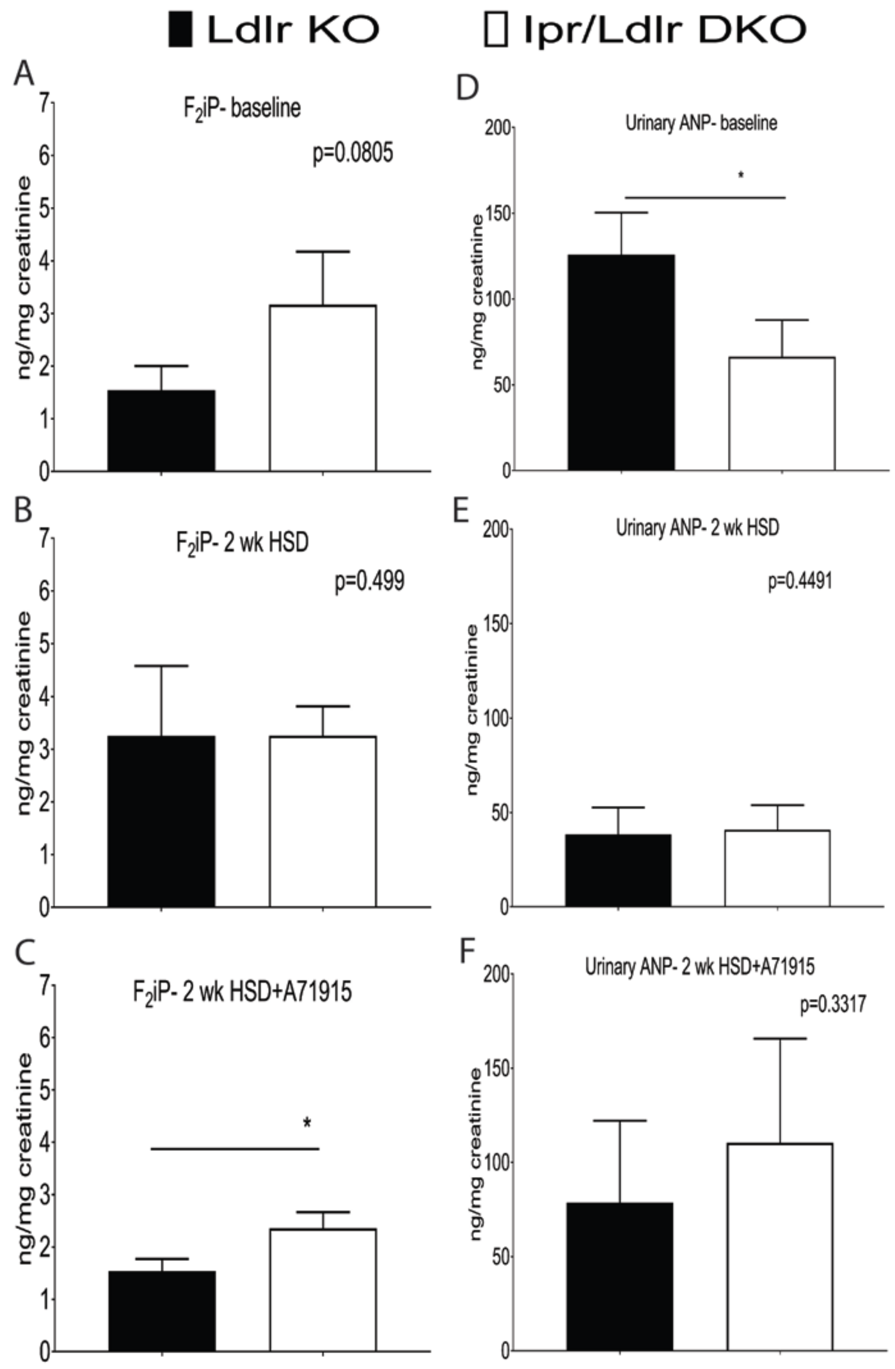


**Supplemental Figure 12. The effect of combined deletion of Ipr and salt-evoked hypertension on urinary $F_2$-isoprostane and atrial natriuretic peptide in female mice.** Urinary $F_2$-isoprostane ($F_2$iP) was analyzed by liquid chromatography/ mass spectrometry as described in the methods. (A) Deletion of Ipr in Ldlr KOs did not significantly increase baseline

(A) and two weeks (B) after an HSD compared with Ldlr KOs (n=11-13). Deletion of Ipr and blockade of ANP receptor with A71915 (50 µg/Kg BW/ day) significantly increase urinary $F_2$iP after feeding an HSD (C, n=5). Urinary ANP was measured using an ELISA kit following manufacturer's instructions. (D) Deletion of Ipr in Ldlr KOs significantly decrease baseline urinary ANP compared with Ldlr KOs (n=11-13). This difference was abolished two weeks after feeding the mice an HSD alone (E) and in conjunction with the ANP receptor antagonist, A71915 (F, n=5). Data are expressed as means ± SEMs (Parametric test, Welch's correction, one-tailed, *$p$< 0.05). We used a one-tailed test for urinary $F_2$iP and ANP because both mediators have been shown to restrain oxidative stress in the vasculature.

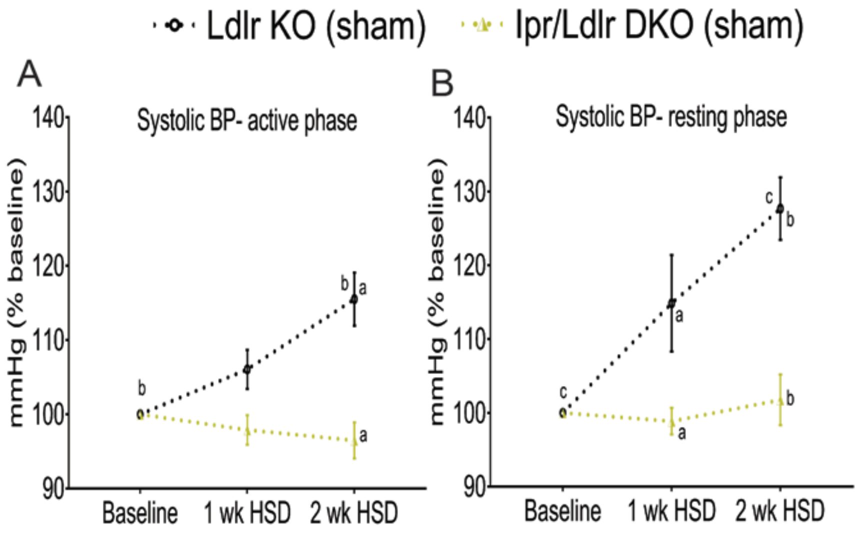


**Supplemental Figure 13. Saline infused via mini-pumps did not rescue hypotension in Ipr-deficient male hyperlipidemic mice fed a high salt diet.** Systolic blood pressures (SBP, A- active phase, B- resting phase) of male mice with mini-pumps containing saline were measured using a tail-cuff system before, one and two weeks after feeding an HSD. Two-way ANOVA revealed a significant effect of genotype and/ or feeding time on SBP responses during the active phase (genotype, $p$< 0.05, feeding time, $p$< 0.0001, interaction, $p$< 0.001) and resting phase (genotype, $p < 0.01$, feeding time, $p < 0.0001$, interaction, $p < 0.01$). Multiple comparison tests (Holm-Sidak) were used to test significant differences between Ldlr KO and Ipr/Ldlr DKO and among different feeding times. Genotype and/ or feeding time with the same lower case letter are significantly different (a- c). Data are expressed as means ± SEMs. n=6 per genotype.

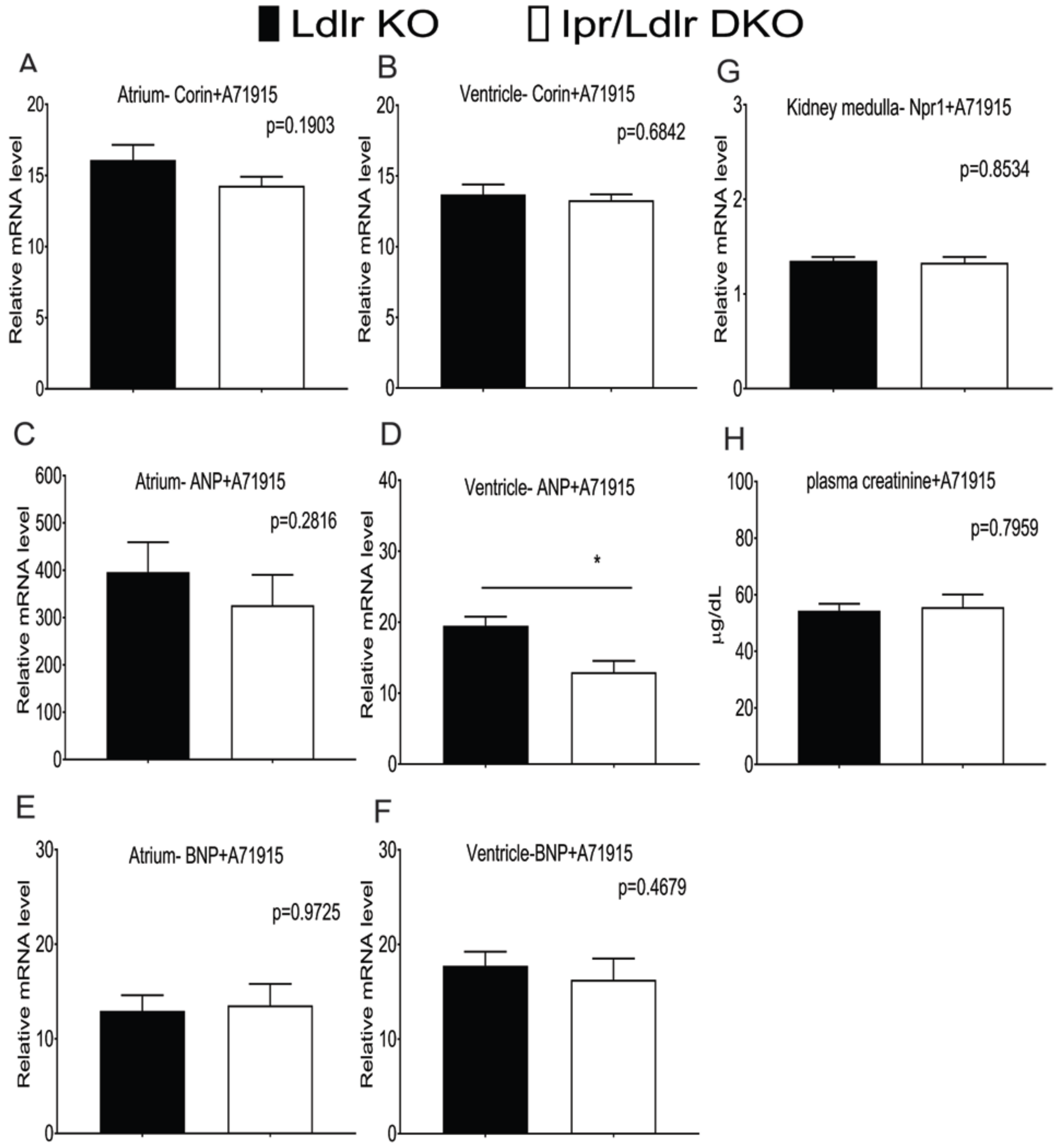

**Supplemental Figure 14. Blockade of the ANP receptor with A71915 abolished the differences in corin, ANP and BNP transcripts in heart atrium and ventricle and Npr1 in kidney medulla between Ipr/Ldlr DKO and Ldlr KO, as measured by RT-qPCR.** Heart atrium, heart ventricle and kidney medulla tissue samples were dissected and RNAs were extracted from Ldlr KO and Ipr/Ldlr DKOs fed an HSD for two weeks in conjunction with ANP inhibition via A71915 infusion (50µg/Kg BW/day) were used for real-time PCR analyses as described in the Methods. Plasma level of creatinine was measured by LC-MS as an indicator of kidney function in mice administered with A71915. No significant differences were detected in corin (A&B), ANP (C), BNP (E&F) and Npr1 (G) transcripts and plasma creatinine levels (H) between Ldlr KO and Ipr/Ldlr DKOs. Ventricle ANP transcript was significantly reduced in Ipr/Ldlr DKO compared with Ldlr KO (D). Data are expressed as means ± SEMs (Parametric test, two-tailed, *$p < 0.05$; n=10 per genotype).

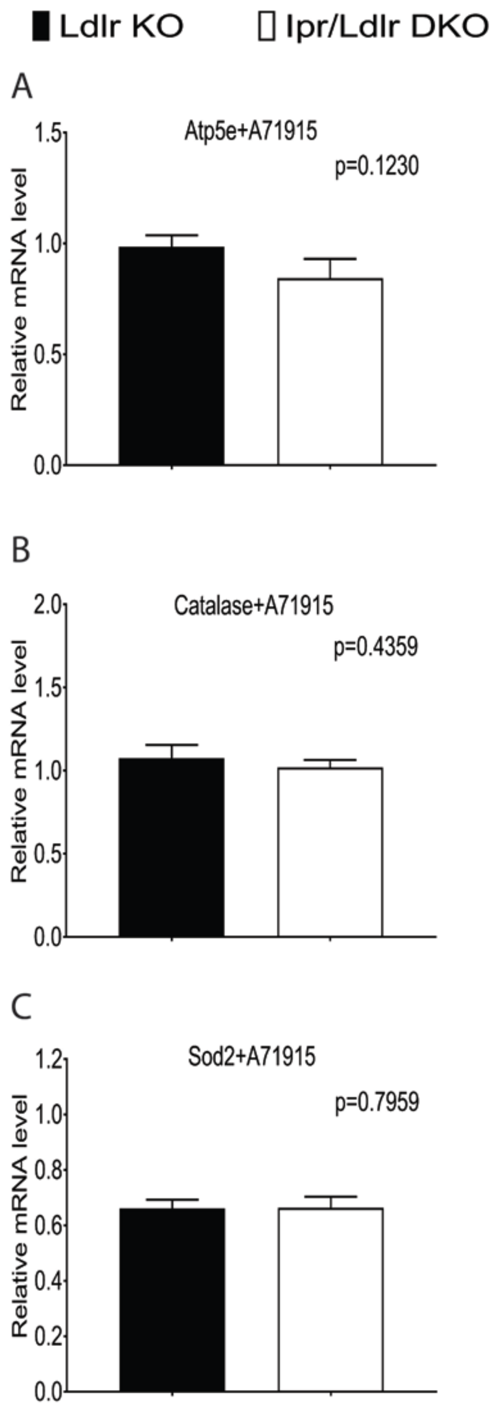


**Supplemental Figure 15. Blockade of the ANP receptor with A71915 abolished the genotype dependent differences in mitochondrial dysfunction and oxidative phosphorylation pathway transcripts in kidney medulla of male mice, as measured by RT-qPCR.** RNA samples isolated from kidney medulla of Ldlr KO and Ipr/Ldlr DKO after two weeks on an HSD in conjunction with ANP inhibition via A71915 infusion (50μg/Kg BW/day) were used for real-time PCR analyses. No significant differences in Atp5e (A), catalase (B) and Sod2 (C) were observed between Ldlr KO and Ipr/Ldlr DKO mice. Data are expressed as means ± SEMs (Parametric test, two-tailed, $p > 0.05$; n=10 per genotype).

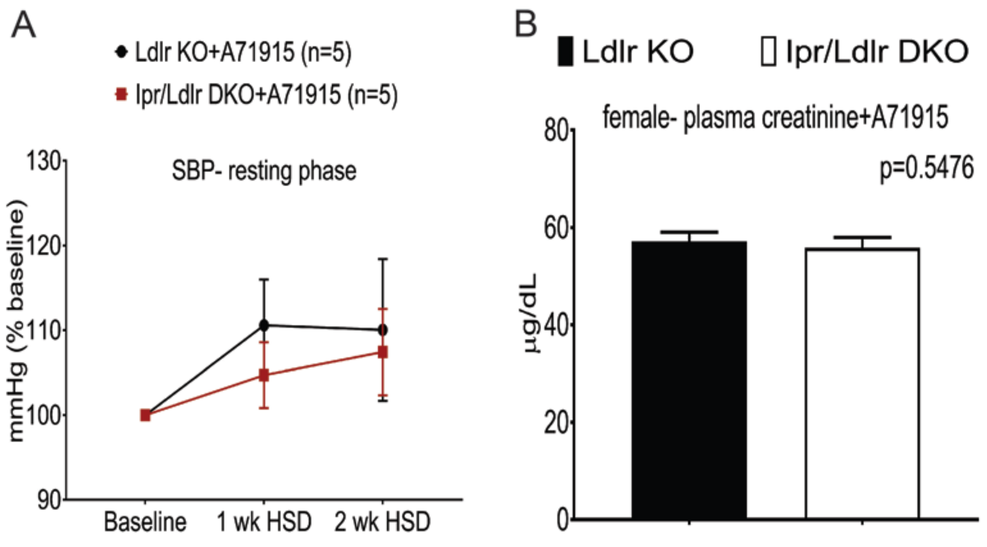


**Supplemental Figure 16. Deletion of Ipr did not alter systolic blood pressure and plasma creatinine in female mice fed a high salt diet in conjunction with an ANP receptor antagonist.** (A) The atrial natriuretic peptide receptor antagonist (A71915, 50μg/Kg BW/day) did not alter SBPs in female Ip-deficient hyperlipidemic mice. A71915 was delivered via mini-pumps and BPs was measured using a tail-cuff system before, one and two weeks after feeding a HSD. Two-way ANOVA revealed no significant effect of feeding time and genotype on SBP in Ldlr KO and Ipr/Ldlr DKO mice. Data are expressed as means ± SEMs. $p > 0.05$; n=5 per genotype. (B) Plasma creatinine level was measured by LC-MS as described in the Methods. Deletion of Ip did not significantly alter plasma creatinine compared with Ldlr KO littermates fed an HSD+ A71915 for two weeks. Data are expressed as means ± SEMs (Parametric test, two-tailed, *p*> 0.05; n=5 per genotype).

**Supplemental Figure 17. Statistical analysis of SBP with an ANP receptor antagonist (A71915) across time in male mice**. 3-way ANOVA with repeated measures and Greenhouse-Geisser sphericity correction showed how SBP changes for different genotypes across time at active/inactive phase, where Ipr and A71915 are the between-subjects factors while week is the repeated measure. ANOVA indicated that SBP is significantly affected by Ipr, A71915 and week and the Ipr:week interaction at active phase, while in the resting phase there is additional significant effect of the A71915:week interaction. The significant factors and interactions are listed below.

Active phase:

- Ipr, F(1,32) = 10.172884, p = 3.182845e-03
- A71975, F(1,32) = 8.701888, p = 5.900875e-03
- week, F(2,64) = 0.9003809, p = 2.062779e-06
- Ipr:week, F(2,64) = 0.9003809, p = 2.885028e-02
- A71975:week, F(2,64) = 0.9003809, p = 3.188379e-02

Resting phase:

- Ipr, F(1,32) = 9.9083548, p = 3.549123e-03
- A71975, F(1,32) = 4.8523003, p = 3.492932e-02
- week, F(2,64) = 0.9043208, p = 1.383358e-07
- Ipr:week, F(2,64) = 0.9043208, p = 2.735816e-02

**Supplemental Figure 18. Statistical analyses of interactions between SBP, genotypes, treatment (week) and phases in ovariectomized female hyperlipidemic mice**. 3-way ANOVA with repeated measures and Greenhouse-Geisser sphericity correction showed how SBP changes for different genotypes across time at active/inactive phase, where Ipr and E2 are the between-subjects factors while week is the repeated measure. ANOVA indicated that SBP is significantly affected by E2, week and the E2:week interaction at both active and resting phase. The significant factors and interactions are listed below.

Active phase:

- E2, $F(1,26) = 8.3392931$, $p = 7.714758e-03$
- week, $F(2,52) = 0.6703226$, $p = 2.350992e-06$
- E2:week, $F(2,52) = 0.6703226$, $p = 1.078514e-02$

Resting phase:

- E2, $F(1,26) = 7.2338891$, $p = 1.232654e-02$
- week, $F(2,52) = 0.6998376$, $p = 0.0000250206$
- E2:week, $F(2,52) = 0.6998376$, $p = 0.0034673737$

**Statistical analyses of interactions between DBP, genotypes, treatment (week) and phases in ovariectomized female hyperlipidemic mice**. 3-way ANOVA with repeated measures and Greenhouse-Geisser sphericity correction showed how DBP changes for different genotypes across time at active/inactive phase, where Ipr and E2 are the between-subjects factors while week is the repeated measure. ANOVA indicated that DBP is significantly affected by week, and the Ipr:E2 and E2:week interactions at both active and resting phase. The significant factors and interactions are listed below.

Active phase:

- Ipr:E2, $F(1,26) = 5.46038554$, $p = 2.742940e-02$
- week, $F(2,52) = 0.6896006$, $p = 1.878692e-05$
- E2:week, $F(2,52) = 0.6896006$, $p = 1.229147e-02$

Resting phase:

- Ipr:E2, $F(1,26) = 5.954118$, $p = 2.180856e-02$
- week, $F(2,52) = 0.6714117$, $p = 4.303491e-05$
- E2:week, $F(2,52) = 0.6714117$, $p = 4.153677e-03$

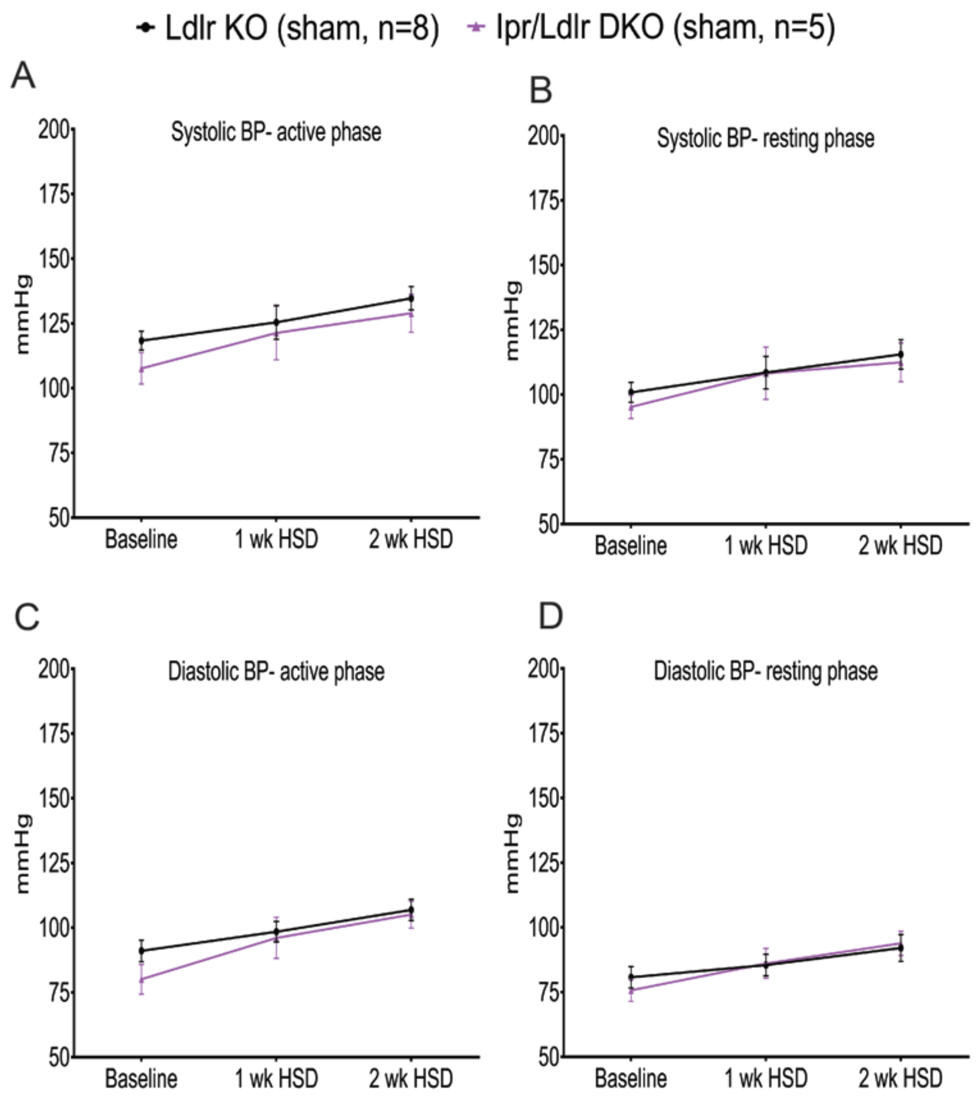


**Supplemental Figure 19. High salt diet failed to alter BP responses in female sham-operated Ldlr KO and Ipr/Ldlr DKO mice.** Salt loading did not significantly alter SBP (A-B) and DBP (C-D) of sham-operated Ipr/Ldlr DKO compared with Ldlr KO mice. Two-way ANOVA revealed no significant effect of genotype and feeding time on SBP and DBP ($p > 0.05$). Data are expressed as means ± SEMs. n=5- 8 per genotype.

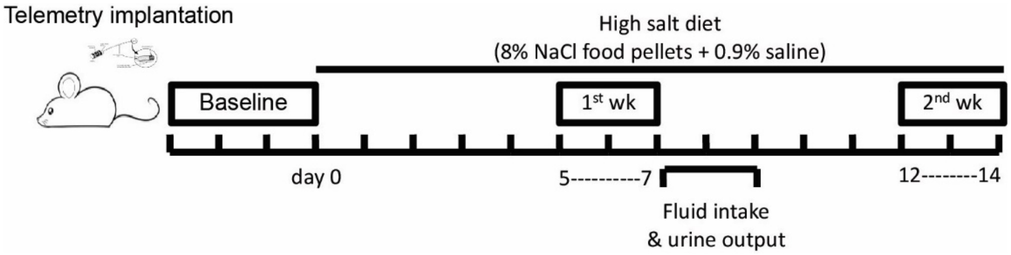


**Supplemental Figure 20. Experimental protocol for the high salt diet regimen.** Eight-to-ten week old mice were implanted with a telemetry transmitter and allowed to recover for at least five days. Baseline blood pressures (SBP and DBP) was recorded for three days before they were placed on a high salt diet (HSD, 8% NaCl+0.9% saline) for two weeks. The recoding system was switched on again from day 5- 7 and day 12-14. Urinary output and fluid intake were estimated by transferring mice into metabolic cages one week after the HSD. Fluid and foods were provided *ad libitum* during the experimental period. At the end of two weeks, mice were sacrificed and plasma and tissues were collected for analyses.
